# Sphingosine 1-phosphate-regulated transcriptomes in heterogenous arterial and lymphatic endothelium of the aorta

**DOI:** 10.1101/802892

**Authors:** Eric Engelbrecht, Michel V. Levesque, Liqun He, Michael Vanlandewijck, Anja Nitzsche, Andrew Kuo, Sasha A. Singh, Masanori Aikawa, Kristina Holton, Richard L Proia, Mari Kono, William T. Pu, Eric Camerer, Christer Betsholtz, Timothy Hla

**Affiliations:** Vascular Biology Program, Boston Children’s Hospital, Deapartment of Surgery, Harvard Medical School, Boston, MA, USA; Department of Immunology, Genetics and Pathology, Rudbeck Laboratory, Uppsala University, Uppsala, Sweden; Karolinska Institutet/AstraZeneca Integrated Cardio Metabolic Centre (KI/AZ ICMC), Karolinska Institutet, Blickagången 6, SE-141 57 Huddinge, Sweden.; Université de Paris, INSERM U970, Paris Cardiovascular Research Center, Paris, France; Center for Interdisciplinary Cardiovascular Sciences, Department of Medicine, Brigham and Women’s Hospital, Harvard Medical School, Boston, MA; Harvard Medical School Research Computing, Boston, MA, USA; Genetics of Development and Disease Branch, National Institute of Diabetes and Digestive and Kidney Diseases, National Institutes of Health, Bethesda, MD, USA; Department of Cardiology, Boston Children’s Hospital, Harvard Medical School, Boston, MA, USA; Harvard Stem Cell Institute, Harvard University, Cambridge, MA, USA

## Abstract

Despite the medical importance of G protein-coupled receptors (GPCRs), *in vivo* cellular heterogeneity of GPCR signaling and downstream transcriptional responses are not understood. We report the comprehensive characterization of transcriptomes (bulk and single-cell) and chromatin domains regulated by sphingosine 1-phosphate receptor-1 (S1PR1) in adult mouse aortic endothelial cells. First, S1PR1 regulates NFkB and nuclear glucocorticoid receptor pathways to suppress inflammation-related mRNAs. Second, spatially distinct S1PR1 signaling in the aorta is associated with heterogenous endothelial cell (EC) subtypes. For example, a transcriptomically distinct arterial EC population at vascular branch points (aEC1) exhibits ligand-independent S1PR1/ß-arrestin coupling. In contrast, circulatory S1P-dependent S1PR1/ß-arrestin coupling was observed in non-branch point aEC2 cells that exhibit an inflammatory signature. Moreover, an adventitial lymphatic EC (LEC) population shows suppression of lymphangiogenic and inflammation-related transcripts in a S1P/S1PR1-dependent manner. These insights add resolution to existing concepts of GPCR signaling and S1P biology.

## Introduction

Sphingosine 1-phosphate (S1P), a circulating lipid mediator, acts on G protein-coupled S1P receptors (S1PRs) to regulate a variety of organ systems. S1PR1, abundantly expressed by vascular endothelial cells (ECs), responds to both circulating and locally-produced S1P to regulate vascular development, endothelial barrier function, vasodilatation and inflammation (Proia & Hla, 2015).

S1P binding to S1PR1 activates heterotrimeric G*_α_*_i/o_ proteins, which regulate downstream signaling molecules such as protein kinases, small GTPases, and other effector molecules to influence cell behaviors, such as shape, migration, adhesion and cell-cell interactions. Even though S1PR1 signaling is thought to evoke transcriptional responses that couple rapid signal transduction events to long-term changes in cell behavior, such mechanisms are poorly understood, especially in the vascular system.

Subsequent to G*_α_*_i/o_ protein activation, the S1PR1 C-terminal tail gets phosphorylated and binds to ß-arrestin, leading to receptor desensitization and endocytosis (Liu et al., 1999; Oo et al., 2007). While S1PR1 can be recycled back to the cell surface for subsequent signaling, sustained receptor internalization brought about by supra-physiological S1P stimulation or functional antagonists leads to recruitment of ubiquitin ligases and lysosomal/proteasomal degradation of the receptor (Oo et al., 2011). Thus, ß-arrestin coupling down-regulates S1PR1 signals. However, studies of other GPCRs suggest that ß-arrestin coupling could lead to biased signaling distinct from G*_α_*_i/o_ -regulated events (Wisler, Rockman, & Lefkowitz, 2018). Distinct transcriptional changes brought about by G*_α_*_i/o_ – and ß-arrestin-dependent pathways are not known.

Our understanding of GPCR signaling *in vivo* is limited. To address this, Kono et al. (2014) developed S1PR1 reporter mice which record receptor activation at single-cell resolution (Kono et al., 2014). Adapted from the “Tango” system (Barnea et al., 2008), S1PR1-GFP signaling (S1PR1-GS) mice convert ß-arrestin recruitment to the GPCR to H2B-GFP expression (Kono et al., 2014). We previously used the S1PR1-GS mouse and showed that high levels of endothelial GFP expression (i.e. S1PR1/ß-arrestin coupling) are prominent at the lesser curvature of the aortic arch and the orifices of intercostal branch points (Galvani et al., 2015). In addition, inflammatory stimuli (e.g. lipopolysaccharide) induced rapid coupling of S1PR1 to ß-arrestin and GFP expression in endothelium in an S1P-dependent manner (Kono et al., 2014). These data suggest that the S1PR1-GS mouse is a valid model to study GPCR activation in vascular ECs *in vivo*.

To gain insights into the molecular mechanisms of S1PR1 regulation of endothelial transcription and the heterogenous nature of S1PR1 signaling *in vivo*, we performed bulk transcriptome and open chromatin profiling of GFP^high^ and GFP^low^ aortic ECs from S1PR1-GS mice. We also performed transcriptome and open chromatin profiling of aortic ECs in which *S1pr1* was genetically ablated (*S1pr1-ECKO*) (Galvani et al., 2015). In addition, we conducted single-cell (sc) RNA-seq of GFP^low^ and GFP^high^ aortic ECs. Our results show that S1PR1 suppresses the expression of inflammation-related mRNAs by inhibiting the NFkB pathway. Second, the high S1PR1 signaling ECs (GFP^high^ cells) are more similar to *S1pr1-ECKO* ECs at the level of the transcriptome. Third, scRNA-seq revealed eight distinct aorta-associated EC populations including six arterial EC subtypes, adventitial lymphatic ECs, and venous ECs, the latter likely from the vasa vasorum. S1PR1 signaling was highly heterogenous within these EC subtypes but was most frequent in adventitial LECs and two arterial EC populations. Immunohistochemical analyses led to anatomical localization of these aortic EC populations and defined markers. In the lymphatic ECs of the aorta, S1PR1 signaling restrains inflammatory and immune-related transcripts. These studies provide a comprehensive resource of transcriptional signatures in aortic ECs, which will be useful to further investigate the multiple roles of S1P in vascular physiology and disease.

## Results

### Profiling the transcriptome of GFP^high^ and GFP^low^ mouse aortic endothelium

S1PR1 expression in aortic endothelium is relatively uniform (Galvani et al., 2015). However, S1PR1 coupling to ß-arrestin, as reported by H2B-GFP expression in S1PR1-GS mice, exhibits clear differences in specific areas of the aorta. For example, thoracic aortae of S1PR1-GS mice show high levels of GFP expression in ECs at intercostal branch points (Galvani et al., 2015), but not in ECs of control mice harboring only the *H2B-GFP* reporter allele (Figure 1A). The first 2-3 rows of cells around the circumference of branch point orifices exhibit the greatest GFP expression (Figure 1A). In addition, heterogeneously dispersed non-branch point GFP+ ECs were also observed, including at the lesser curvature of the aortic arch (Figure 1A). Areas of the aorta that are distal (> ∼10 cells) from branch points, as well as the greater curvature, exhibit relatively low frequencies of GFP+ ECs (Figure 1A). GFP+ mouse aortic ECs (MAECs) are not co-localized with Ki-67, a marker of proliferation, suggesting that these cells are not actively cycling (Figure 1A). However, fibrinogen staining was frequently co-localized with GFP+ MAECs, suggesting that ß-arrestin recruitment to S1PR1 was associated with increased vascular leak (Figure 1A). These findings suggest sharp differences in S1PR1 signaling throughout the normal mouse aortic endothelium.

**Figure 1.**
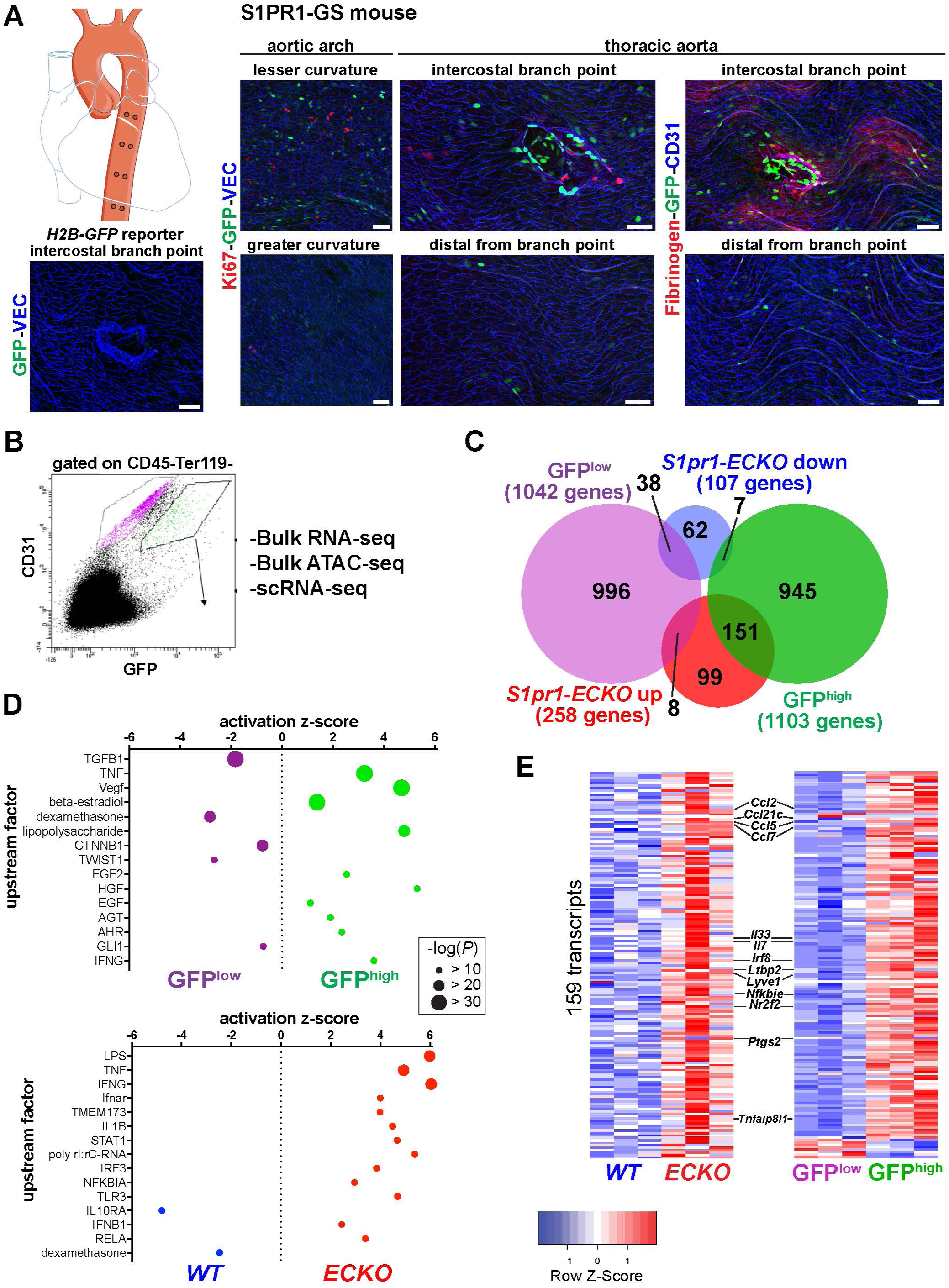
High S1PR1/ß-arrestin coupling in normal mouse aortic endothelium exhibits transcriptomic concordance with S1PR1 loss-of-function. (A) *H2B-GFP* control and S1PR1-GS mouse thoracic aorta whole mount *en face* preparations. Representative images from different regions of the aorta are presented and show H2B-GFP (GFP), VE-Cadherin (VEC) or CD31, Ki67 (N=3) or Fibrinogen (N=2) immunostaining. Scale bars are 50 µM. **(B)** FACS gating scheme used for isolation of GFP^high^ and GFP^low^ MAECs. *S1pr1-ECKO* and *WT* MAECs were isolated using the GFP^low^ gate of this scheme (see Figure 1-figure supplement 1A). **(C)** Venn diagram showing differentially expressed transcripts in the GFP^high^ vs. GFP^low^ and *S1pr1-ECKO* vs. *WT* MAECs comparisons. The number of transcripts individually or co-enriched (p-value < 0.05) are indicated for each overlap. **(D)** Selected upstream factors identified by IPA analysis of GFP^high^ vs. GFP^low^ (top) and *S1pr1-ECKO* vs. *WT* (bottom) MAEC comparisons. Activation Z-scores and P-values are indicated for each selected factor. **(E)** Expression heatmaps (row Z-scores) of the 159 *S1pr1-ECKO* up-regulated transcripts also differentially expressed between GFP^high^ and GFP^low^ MAECs. Selected transcripts are labeled between the heatmaps from the comparison of *S1pr1-ECKO* vs *WT* (left) and GFP^high^ vs GFP^low^ (right).

For insight into the aortic endothelial transcriptomic signature associated with high levels of S1PR1/ß-arrestin coupling, we harvested RNA from fluorescent-activated cell sorted (FACS) GFP^high^ and GFP^low^ MAECs and performed RNA-seq (Figure 1B). To identify genes that are regulated by S1PR1 signaling, we sorted MAECs from tamoxifen-treated *Cdh5*-Cre^ERT2^ *S1pr1*^f/f^ (*S1pr1*-*ECKO*) and *S1pr1*^f/f^ (*S1pr1*-*WT*) littermates (Figure 1-figure supplement 1A). We noted that GFP^high^, GFP^low^, *S1pr1*-*WT* and *S1pr1*-*ECKO* MAECs each expressed endothelial lineage genes (*Pecam1*, *Cdh5*) and lacked hematopoietic and VSMC markers (*Ptprc, Gata1,* and *Myocd*), validating our MAEC isolation procedure (Figure 1-figure supplement 1B). Efficient CRE-mediated recombination of *S1pr1* was confirmed in sorted MAECs from *S1pr1*-*ECKO* mice (Figure 1-figure supplement 1B).

Differential expression analysis using edgeR identified 1,103 GFP^high^-enriched and 1,042 GFP^low^-enriched transcripts (p-value < 0.05) (Figure 1C and Figure 1-figure supplement 1B; see also Supplementary File 1). In contrast, *S1pr1*-*ECKO* MAECs showed fewer differentially expressed genes (DEGs), with 258 up- and 107 down-regulated transcripts (Figure 1C and Figure 1-figure supplement 1C; see also Supplementary File 1). Intersection of these two sets of DEGs showed that only 9.5% (204 transcripts) were common (Figure 1C and Supplementary File 1), suggesting that the majority (∼90%) of transcripts that are differentially expressed in MAECs from S1PR1-GS mice are not regulated by S1PR1 signaling. Rather, S1PR1/ß-arrestin coupling may correlate with heterogenous EC subtypes in the mouse aorta.

Among the 204 common DEGs, 151 were both *S1pr1*-*ECKO* upregulated and enriched in the GFP^high^ population (Figure 1C). In contrast, much lower numbers of transcripts were found in the intersection of GFP^high^ and *S1pr1*-*ECKO* downregulated (7 transcripts), GFP^low^ and *S1pr1*- *ECKO* upregulated (8 transcripts) and GFP^low^ and *S1pr1*-*ECKO* downregulated (38 transcripts) (Figure 1C). We computed the statistical significance of these gene set overlaps using the GeneOverlap R package (Shen et al., 2013). The GFP^high^:*S1pr1*-*ECKO*-up overlap was significant (151 transcripts, p-value = 3.80E-126), as was the GFP^low^:*S1pr1*-*ECKO*-down overlap (38 transcripts, p-value = 3.80E-126), and the other two tested overlaps were not significant (p-value > 0.3) (Figure 1-figure supplement 1E). These data suggest that the transcriptome of MAECs exhibiting high S1PR1/ß-arrestin coupling (GFP^high^) is more similar to that of *S1pr1*-*ECKO*.

We used Ingenuity Pathway Analysis (IPA®, Qiagen) to examine biological processes regulated by S1PR1 signaling and loss of function in MAECs. Transcripts involved in inflammatory processes were prominently up-regulated in both GFP^high^ and *S1pr1*-*ECKO* MAECs (Figure 1D; see also Supplementary File 2). For example, positive tumor necrosis factor (TNF)-a, lipopolysaccharide, and interferon-g signaling were observed in both *S1pr1*-*ECKO* and GFP^high^ MAECs. In contrast, a negative glucocorticoid signature was observed in these cells (Figure 1D). Examples of differentially regulated transcripts are chemokines (*Ccl2*, *Clc5*, *Ccl7, Ccl21c*), cytokines (*Il33*, *Il7*), inflammatory modulators (*Irf8*, *NF*k*Bie, TNFaip8l1*) and cyclooxygenase-2 (*Ptgs2*) (Figure 1E and Figure 1-figure supplements 1C and 1D). This suggests that S1PR1 suppresses inflammatory gene expression in mouse aortic endothelium. We noted that transcripts in the TGFß signaling pathway (*Thbs1*, *Smad3*, *Bmpr1a*, *Col4a4*, *Pcolce2*) were prominently downregulated in the GFP^high^ population (Figure 1D and Figure 1-figure supplement 1C). Furthermore, both GFP^high^ and *S1pr1-ECKO* MAECs were enriched with *Lyve1*, *Flt4*, and *Ccl21c* transcripts, which encode proteins with well-defined roles in lymphatic EC (LEC) differentiation and function (Ulvmar & Makinen, 2016) (Figure 1E, Figure 1-figure supplement 1F). Taken together, these data suggest that S1PR1 represses expression of inflammatory genes in aortic endothelium and that GFP^high^ MAECs include aorta-associated LECs and are heterogeneous. **Chromatin accessibility landscape of MAECs**

We used the assay for transposase-accessible chromatin with sequencing (ATAC-seq) (Buenrostro, Giresi, Zaba, Chang, & Greenleaf, 2013) to identify putative *cis*-elements that regulate differential gene expression between GFP^high^ versus GFP^low^ and *S1pr1-WT* versus *S1pr1- ECKO* MAECs. ATAC-seq utilizes a hyper-active Tn5 transposase (Adey et al., 2010) that simultaneously cuts DNA and ligates adapters into sterically unhindered chromatin. This allows for amplification and sequencing of open chromatin regions containing transcriptional regulatory domains such as promoters and enhancers. After alignment, reads from three experiments were trimmed to 10 bp, centered on Tn5 cut sites, then merged. These merged reads were used as inputs to generate two peak sets (MACS2, FDR < 0.00001) of 73,492 for GFP^low^ MAECs and 65,694 for GFP^high^ MAECs (Figure 2A). MAECs isolated from *WT* and *S1pr1*-*ECKO* mice harbored 93,859 and 76,082 peaks, respectively (Figure 2A). Peaks were enriched in promoter and intragenic regions (Figure 2-figure supplement 1A). We noted that the *Cdh5* gene exhibited numerous open chromatin peaks, while *Gata1* was inaccessible (Figure 2-figure supplement 1B). Furthermore, we observed a global correlation between chromatin accessibility and mRNA expression for all 20,626 annotated coding sequences (CDS’s) in the NCBI RefSeq database (Figure 2-figure supplement 2). These data suggest that our ATAC-seq data is of sufficient quality for detailed interrogation.

**Figure 2.**
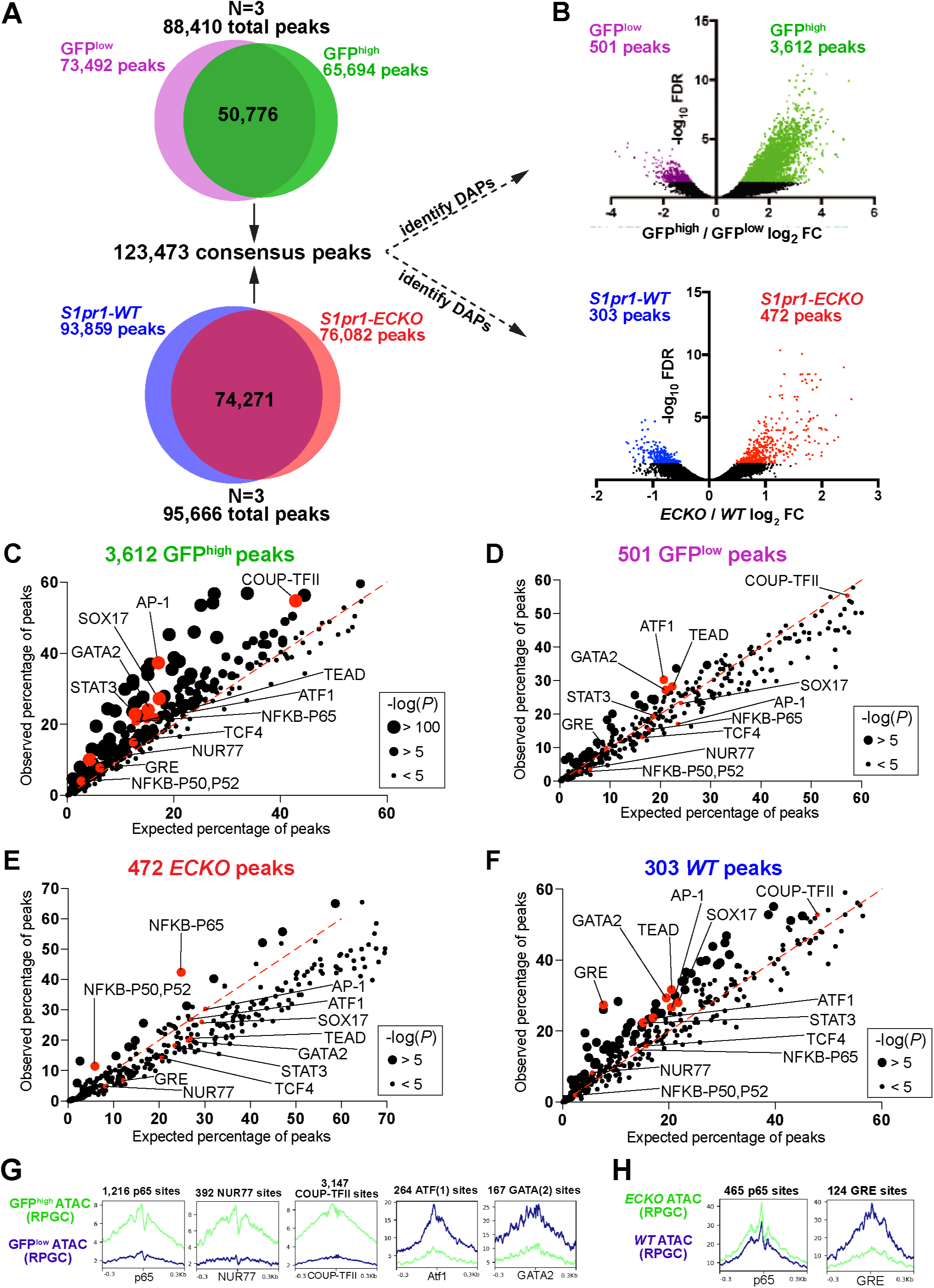
**S1PR1 loss-of-function and high levels of ß-arrestin coupling are associated with an NF**k**B signature in open chromatin. (A)** Venn diagrams illustrating all peaks (FDR < 0.00001) identified after analysis of individual ATAC-seq replicates of GFP^low^, GFP^high^ *S1pr1-ECKO*, and *WT* MAECs, and subsequent merging of these peaks into a single consensus peak set. **(B)** Three individual experiments were performed for GFP^low^ vs GFP^high^ and *S1pr1-ECKO* vs *WT* comparisons. Differentially accessible peaks (DAPs) were determined using edgeR (FDR < 0.05, see Methods). **(C-F)** DAPs were input to the HOMER “findMotifsGenome.pl” script (see also Supplementary File 2) and observed vs expected frequencies of motif occurrances were plotted. **(G-H)** ATAC-seq reads were centered on Tn5 cut sites, trimmed to 10 bp, and nucleotide-resolution bigwig files were generated using DeepTools with reads per genomic content (RPGC) normalization. Reads were subsequently centered on TF binding motifs identified in **(C-F)** and viewed as mean read densities across 600 bp windows.

Differential chromatin accessibility analysis of GFP^high^ versus GFP^low^ MAECs identified 501 peaks with reduced accessibility (GFP^low^ peaks) and 3,612 peaks with greater accessibility (GFP^high^ peaks) in GFP^high^ MAECs (FDR < 0.05, Figure 2B and Supplementary File 3). For *WT* and *ECKO* counterparts, this analysis identified 303 peaks with reduced accessibility (*S1pr1-WT* peaks) and 472 peaks with enhanced accessibility (*S1pr1-ECKO* peaks) in *S1pr1-ECKO* MAECs (Figure 2B and Supplementary File 3). The ∼7-fold higher number of GFP^high^-enriched peaks suggests that GFP^high^ MAECs are more “activated” (elevated number of chromatin remodeling events) and/or are a heterogeneous mixture of EC subtypes.

To identify relevant transcription factors (TFs), we used the Hypergeometric Optimization of Motif EnRichment (HOMER) (Heinz et al., 2010) suite of tools to reveal over-represented motifs in each set of differentially accessible peaks (DAPs). GFP^high^ peaks were enriched with p65-NFkB, AP-1, STAT3, SOX17, COUP-TFII, and NUR77 motifs (Figure 2C and Supplementary File 3). In contrast, GFP^low^ peaks showed reduced occurrence of these motifs (Figure 2D). *S1pr1*-*ECKO* peaks were enriched with p65-NFkB motifs, while *S1pr1*-*WT* peaks were markedly enriched with glucocorticoid response elements (GREs) and modestly enriched with STAT3, GATA2, ATF1, SOX17 and COUP-TFII motifs (Figure 2E and 2F). Examination of ATAC-seq reads centered on selected binding sites (p65, NUR77, COUP-TFII, ATF1, GATA2, and GRE) showed local decreases in accessibility at motif centers, suggestive of chromatin occupancy by these factors (Figures 2G and 2H).

We used the ATAC-seq footprinting software HINT-ATAC (Z. Li et al., 2019) to assess genome-wide putative chromatin occupancy by TFs. HINT-ATAC identified enhanced footprinting scores at NFKB1 and NFKB2 motifs in *S1pr1*-*ECKO* MAECs, whereas GFP^high^ MAECs showed increasesd scores at RELA motifs and to a lesser extent at NFKB1 and NFKB2 motifs (Figure 2-figure supplement 3A and 3B). This analysis also identified motifs of the TCF/LEF family (LEF1, TCF7, TCF7L2) as GFP^high^-enriched, but not *S1pr1*-*ECKO*-enriched (Figure 2-figure supplement 3A-D). Consistent with HOMER analysis of DAPs, HINT-ATAC identified NR2F2 (COUP-TFII), NR4A1 (NUR77), TCF4, and SOX17 motifs as exhibiting greater footprinting scores in GFP^high^ MAECs, while GFP^low^ MAECs showed enhanced putative chromatin occupancy at ATF1 motifs.

**Figure 3.**
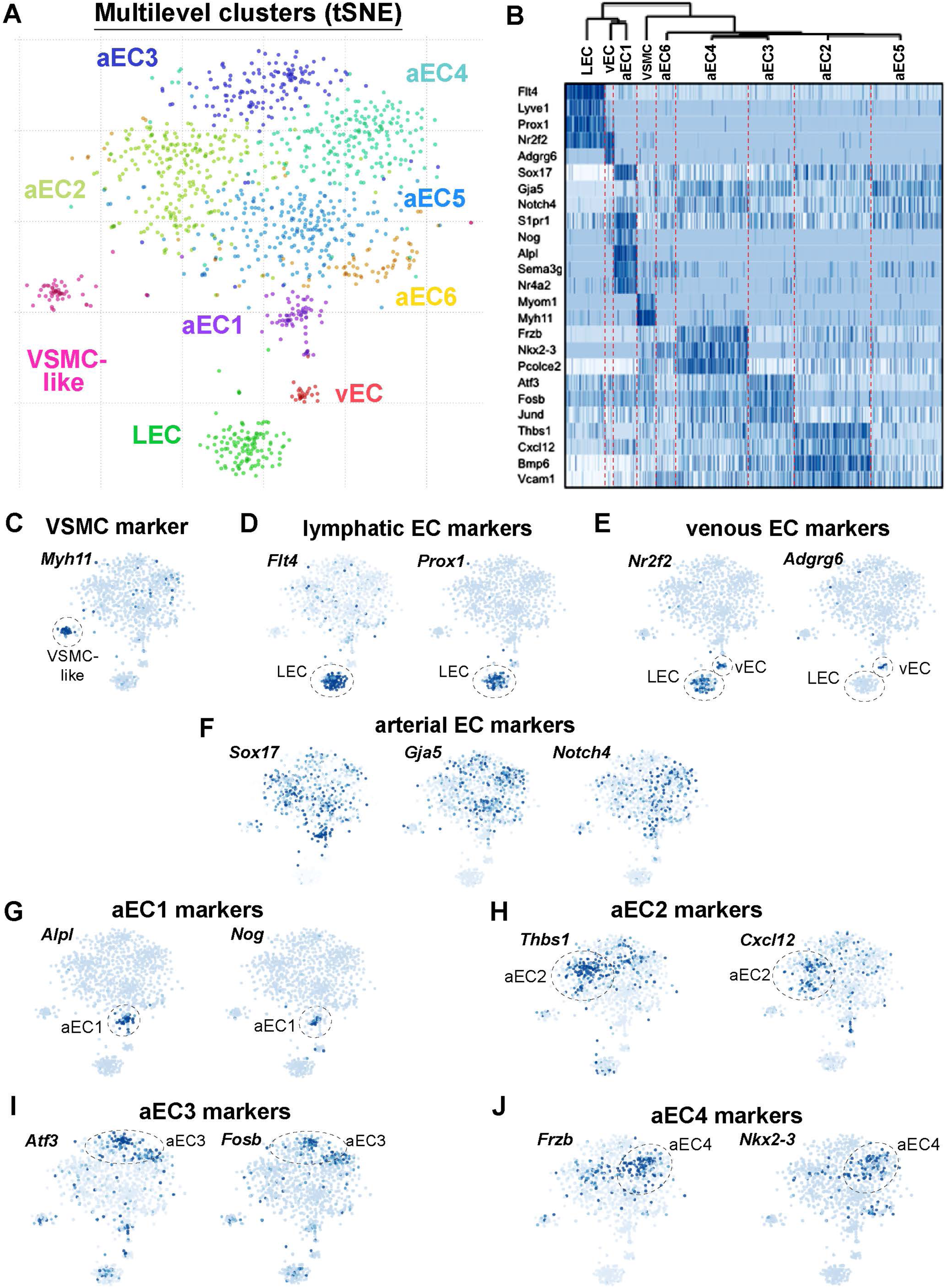
Single-cell RNA-sequencing of GFP^high^ and GFP^low^ MAECs. (A) t-SNE projection from Pagoda2 multilevel clustering of 767 GFP^high^ and 384 GFP^low^ MAECs. Dash-line circles highlight each of the 9 clusters identified. Cells and cluster names are color- coded according to cluster assignment. **(B)** Dendrogram from hierarchical clustering and expression heatmap of selected genes. The dendrogram (top) reveals an upstream split between LEC, vEC, aEC1 and aEC2, aEC3, aEC4, aEC5, and aEC6 populations. The heatmap shows the gradient of expression, from low (white) to high (dark blue), for a selection of transcripts with distinctive expression patterns. **(C-F)** Expression of transcripts specific to VSMCs **(C)** lymphatic ECs **(D)**, venous ECs **(E)** and arterial ECs **(F)** are shown on the t-SNE embedding. **(G-J)** Representative transcripts enriched in arterial EC (aEC) clusters 1 (G), 2 (H), 3 (I) and 4 (J) are shown on the t-SNE embedding.

Analysis of DAPs showed that only the p65-NFkB motif was commonly enriched between GFP^high^ and *S1pr1*-*ECKO* MAECs. This observation is consistent with our RNA-seq analysis, which identified cytokine/NFkB pathway suppression by S1PR1 signaling in MAECs (Figure 1D). Enrichment of COUP-TFII, NUR77, and AP-1/bZIP motifs in open chromatin of GFP^high^ MAECs, but not *S1pr1-ECKO* MAECs, further suggests that high levels of S1PR1/ß-arrestin coupling occurs in heterogenous populations of aortic ECs.

### Single-cell RNA-seq analysis of GFP^high^ and GFP^low^ MAECs reveals 8 distinct EC clusters

Imaging studies demonstrated that GFP^high^ MAECs are restricted to specific anatomical locations. To test the hypothesis that these represent specific EC subpopulations, we employed single-cell RNA-seq (scRNA-seq) on FACS-sorted GFP^high^ and GFP^low^ MAECs. In total, 1152 cells were sequenced (768 GFP^high^ and 384 GFP^low^) using the Smart-seq2 protocol. An average of 300,000 aligned reads/cell were obtained and corresponded to ∼3,200 transcripts/cell. *Cdh5* transcripts were broadly detected, consistent with endothelial enrichment of sorted cells (Figure 3- figure supplement 1A). *S1pr1* and *Arrb2* were also broadly detected (Figure 3-figure supplement 1A), suggesting that receptor activation rather than expression of these factors accounts for heterogenous reporter expression in MAECs.

Analysis of GFP^high^ and GFP^low^ MAECs using a custom velocyto/pagoda2 pipeline (Fan et al., 2016; La Manno et al., 2018) revealed 9 clusters upon T-distributed stochastic neighborhood embedding (t-SNE) projection (Figure 3A). 6 of the 9 clusters grouped together in a “cloud”, whereas 3 clusters formed distinct populations. We used hierarchical differential expression analysis to identify signature marker genes of each cluster (Figure 3B).

Genes uniquely detected in one of the distinct clusters included vascular smooth muscle cell (VSMC)-specific transcripts such as *Myh11*, *Myom1*, and *Myocd* (Figure 3B and 3C; Figure 3-figure supplement 1B). Therefore, this cluster was designated VSMC-like as these cells may represent MAECs sorted along with fragments of VSMCs, or “doublets” of ECs and VSMCs. Because these cells may represent contamination in an otherwise pure pool of aortic ECs, we omitted this VSMC-like cluster from subsequent analyses. The remaining eight EC clusters were further analyzed.

Lymphatic EC (LEC) markers such as *Flt4* (VEGFR3), *Prox1*, and *Lyve1*, as well as the venous marker *Nr2f2* (COUP-TFII), were detected in a distinct cluster (Figures 3B and 3D; Figure 3-figure supplement 1B). A smaller but nonetheless distinct cluster of ECs was also enriched with *Nr2f2* transcripts but lacked lymphatic markers, suggesting that these cells are of venous origin (Figures 3B and 3E; Figure 3-figure supplement 1B). Arterial lineage markers *Sox17, Gja5* and *Notch4* were expressed in the 6 clusters comprising the “cloud” of ECs (aEC1-6) (Figures 3B and 3F; Figure 3-figure supplement 1B).

We individually compared LECs, vECs, and VSMCs to the remainder of ECs as a “pseudo- bulk” cluster to generate a list of transcripts enriched (Z-score > 3) for each of these three clusters. We performed the same analysis for aEC1, aEC2, aEC3, aEC4, aEC5, and aEC6, but used only arterial ECs as the comparator. For example, aEC1-enriched transcripts were identified by generating a “pseudo-bulk” merge of all aEC2, aEC3, aEC4, aEC5, and aEC6 cells, while aEC2- enriched transcrips were compared to the pseudo-bulk merge of aEC1, aEC3, aEC4, aEC5, and aEC6. The top 32 transcripts that resulted from this analysis are shown in Figure 3-figure supplement 2. Among the arterial clusters, aEC1 and aEC2 harbored the greatest numbers of enriched transcripts (Z-score > 3) with 411 and 1517, respectively (Figure 3-figure supplement 3A; see also Supplementary File 4). We noted that aEC5 exhibited the fewest (77) enriched transcripts (Figure 3-figure supplement 3A). Representative marker genes of aEC1, aEC2, aEC3, and aEC4 are shown in t-SNE embedding in Figure 3G-J.

LEC (97% GFP^high^), vEC (100% GFP^high^), aEC1 (97% GFP^high^) and aEC2 (92% GFP^high^) harbored the greatest proportion of GFP^high^ MAECs, suggesting that S1PR1/ß-arrestin coupling is robust in these ECs subtypes (Figure 3-figure supplement 3A and 3B). In contrast, aEC3-6 contained lower frequencies of GFP^high^ MAECs. aEC4 (30% GFP^high^) exhibited the lowest frequency of MAECs with S1PR1/ß-arrestin coupling. The GFP^high^-dominated clusters (LEC, vEC, aEC1, and aEC2) were enriched with several transcripts related to sphingolipid metabolism, such as *Spns2*, *Sptlc2*, *Ugcg*, *Enpp2*, *Ormdl3*, *Degs1*, and *Sgms2* (Figure 3-figure supplement 4A; see also Supplementary File 5). Notably, *S1pr1* transcripts were enriched in aEC1 cells by ∼1.8- fold relative to the remainder of arterial ECs (Figure 3-figure supplement 4A; see also Supplementary File 4).

**Figure 4.**
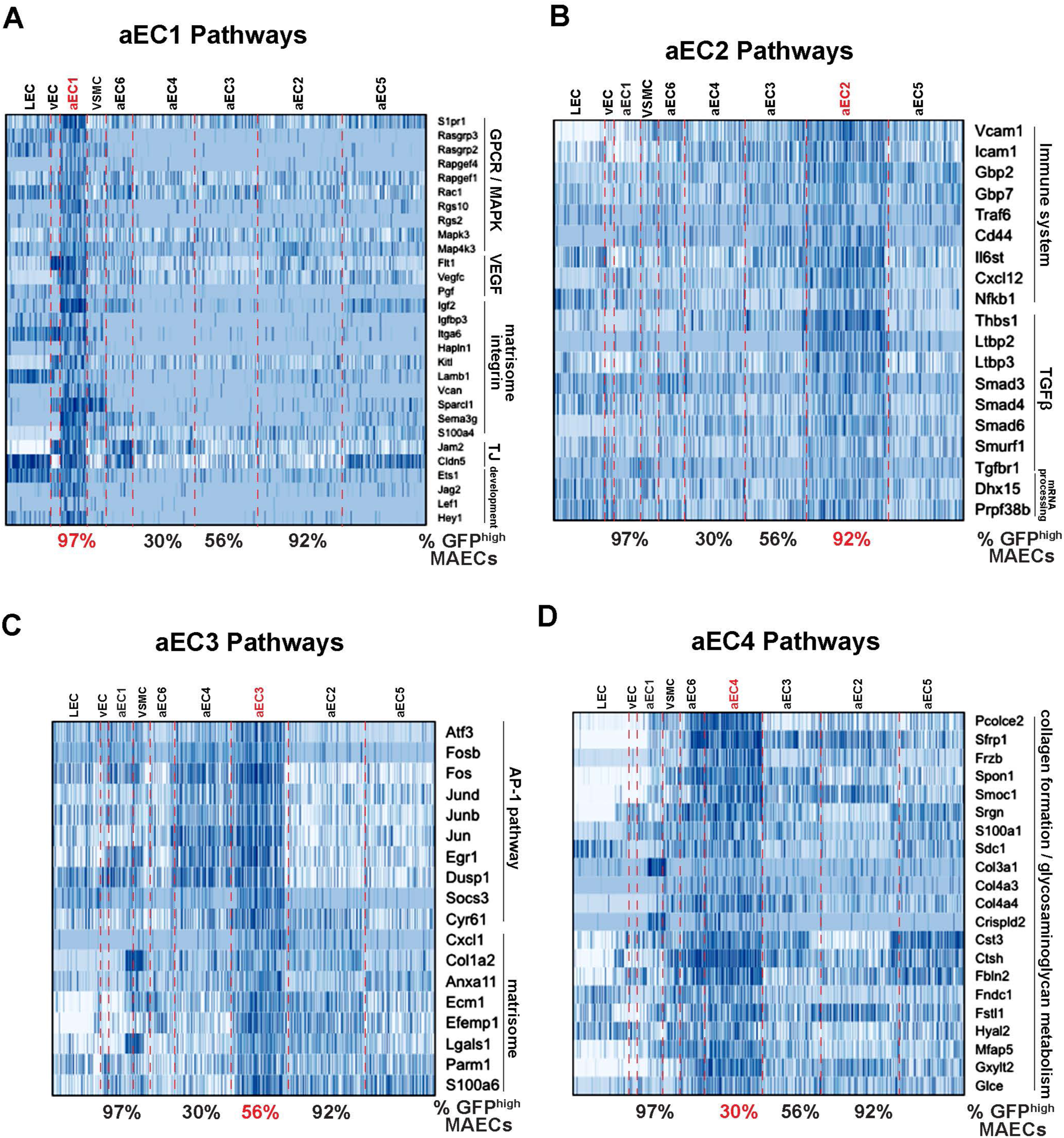
Functional annotation of arterial MAECs clusters aEC1, aEC2, aEC3, and aEC4. (A-D) Selected pathways from GSEA analysis of cluster-enriched transcripts. Pathways enriched in aEC1 **(A)**, aEC2 **(B)**, aEC3 **(C)**, and aEC4 **(D)** are shown with representeative transcripts identified in each pathway (see also Supplementary File 5). The percent of GFP^high^ cells in each of the four analyzed clusters are indicated at the bottom each heatmap.

Pagoda2 clustering suggested that aEC1 cells were more similar to LECs and vECs than to the remainder of arterial ECs. This is illustrated by the first split of the hierarchical clustering dendrogram, which separated LEC, vEC, and aEC1 from the remainder of arterial ECs (aEC2-6) (Figure 3B). To identify genes that underlie the similarity between LEC, vEC, and aEC1, we identified all transcripts commonly enriched (46 transcripts, Z-score > 3) and depleted (92 transcripts, Z-score < -3) in each of these 3 clusters when individually compared to a pseudo-bulk merge of aEC2-6 (Supplementary File 4). Examples of LEC, vEC, and aEC1 co-enriched transcripts were *Itga6*, *Apold1*, *Kdr*, *Fabp4*, *Robo4*, *Tcf4*, and *Adgrf5* (Figure 3-figure supplement 4B). Conversely, *Sod3*, *Pcolce2*, *Col4a4*, *Frzb*, *Sfrp1*, *Gxylt2*, and *Bmpr1a* were depleted from LEC, vEC, and aEC1. Notably, these depleted transcripts were highly enriched in aEC4, which is 30% GFP^high^ MAECs. In contrast, LEC, vEC, and aEC1 are each > 95% GFP^high^.

We note that the abovementioned transcripts (Figure 3-figure supplement 4B) were among those most differentially expressed between GFP^high^ and GFP^low^ MAECs by bulk RNA-seq (Figure 1-figure supplement 1C). This is demonstrative of consistency between our bulk and single-cell datasets.

For functional insights into arterial EC clusters, we analyzed aEC1-aEC4 enriched transcripts with the Gene Set Enrichment Analysis (GSEA) tool (Figures 4A-D). aEC1 cells were enriched with transcripts associated with GPCR/MAPK signaling (*Rasgrp3*, *Rapgef4*, *Rgs10*, *Mapk4k3*, *S1pr1*) as well as VEGF, integrin, and tight-junction pathways (*Flt1*, *Vegfc*, *Pgf*, *Igf2*, *Vcan*, *Sema3g*, *S100a4*, *Jam2*, *Cldn5*). The aEC2 cluster presented a different profile with enriched terms related to immune/inflammatory pathways, TGFß signaling and mRNA processing. Elevated expression of *Vcam1*, *Icam1*, *Traf6*, *Cxcl12* and *NFkb1* may suggest that these ECs represent an inflammatory cluster.

In contrast, aEC3 cells were enriched with “immediate-early” transcripts, including those of the AP-1 transcription factor family (*Atf3*, *Jun*, *Jund*, *Junb*, *Fos*, *Fosb*). Enhanced expression of *Atf3* and related TFs of the bZIP family in aEC3 may have contributed to increased accessibility at ATF, FOSB::JUN, and FOSB::JUNB binding sites in GFP^low^ MAECs (Figure 2D and 2G; Figure 2-figure supplement 3C). Notably, a recent study of young (8-week) and aged (18-month) normal mouse aortic endothelium also identified a cluster of *Atf3*-positive cells only in young endothelium, as determined by both scRNA-seq and immunostaining for ATF3 (McDonald et al., 2018). Markers of these cells were also identified as top aEC3 markers (e.g. *Fosb*, *Jun*, *Jund*, *Junb*, *Dusp1*) suggesting that aEC3 cells are similar to the regenerative *Atf3*-positive cluster described by McDonald et al. (2018).

aEC4 cells were enriched with transcripts related to cell-ECM interations, glycosaminoglycan metabolism, and collagen formation (*Pcolce2*, *Frzb*, *Spon1*, *Col4a4*, *Mfap5*, *Hyal2*). The other two arterial clusters (aEC5 and aEC6) appeared less distinctive (i.e. harbored relatively few enriched transcripts with high Z-scores and fold change values) and therefore were not analyzed. Collectively, these data suggest that scRNA-seq identified more than four distinct MAEC subtypes with unique transcriptomic signatures.

Next, we integrated our dataset with information from two recent scRNA-seq studies that also sub-categorized ECs of the normal mouse aorta (Kalluri et al., 2019; Lukowski et al., 2019). Lukowski et al. (2019) sequenced individual FACS-sorted Lineage^-^CD34^+^ cells and identified 2 major EC clusters, designated “Cluster 1” and “Cluster 2”. Kalluri et al. (2019) identified 3 major EC clusters by sequencing individual cells from whole-aorta digests. Both studies used 10x Genomics droplet sequencing for cell capture and gene expression library preparation. Top markers of “Cluster 1” (Lukowski et al., 2019) were primarily expressed by LEC, vEC and aEC1 (Figure 4-figure supplement 1A). “Cluster 1” shared markers with “EC2” (Kalluri et al., 2019), such as *Rgcc*, *Rbp7*, *Cd36*, *Gpihbp1* (Figure 4-figure supplement 1A and 1B). Similarly, several “EC2” markers, while enriched in aEC1 relative to aEC2-6 (e.g. *Flt1*, *Rgcc*), were also expressed in LEC and vEC (e.g. *Pparg*, *Cd36*, *Gpihbp1, Rbp7*) (Figure 4-figure supplement 1B). “EC3” (Kalluri et al., 2019) markers were strongly enriched in LEC and vEC (e.g. *Nr2f2*, *Flt4*, *Lyve1*), and the authors noted that these cells were likely of lymphatic origin (Figure 4-figure supplement 1B). The remaining two clusters, “Cluster 2” (Lukowski et al., 2019) and “EC1” (Kalluri et al., 2019), strongly resembled the aggregate of aEC2-6 as they were enriched with *Gata6*, *Vcam1*, *Dcn*, *Mfap5*, *Sfrp1*, *Eln*, and *Cytl1* (Figure 4-figure supplement 1A and 1B). Taken together, these information from three independent groups strongly suggest that a major source of heterogeneity in the normal adult mouse aorta, as revealed by scRNA-seq analysis, includes differences between LEC, vEC, aEC1 and aEC2-6.

### Localization of heterogenous arterial EC populations

Among the arterial populations, aEC1 and aEC2 exhibited the highest frequencies of S1PR1/ß-arrestin coupling (> 90%). Thus, we addressed the anatomical location of these arterial clusters in the normal adult mouse aorta by immunolocalization of markers. We utilized antibodies against noggin (NOG), alkaline phosphatase (ALPL), and integrin alpha-6 (ITGA6), which are highly enriched in aEC1, as well as fibroblast-specific protein-1 (FSP1, encoded by *S100a4*) and claudin-5 (CLDN5), which are enriched in, but not exclusive to, aEC1 (Figure 5A). We also utilized antibodies against thrombospondin-1 (TSP1, encoded by *Thbs1*) and vascular cell adhesion molecule 1 (VCAM1), which are enriched in aEC2 and depleted in aEC1 (Figure 5A).

**Figure 5.**
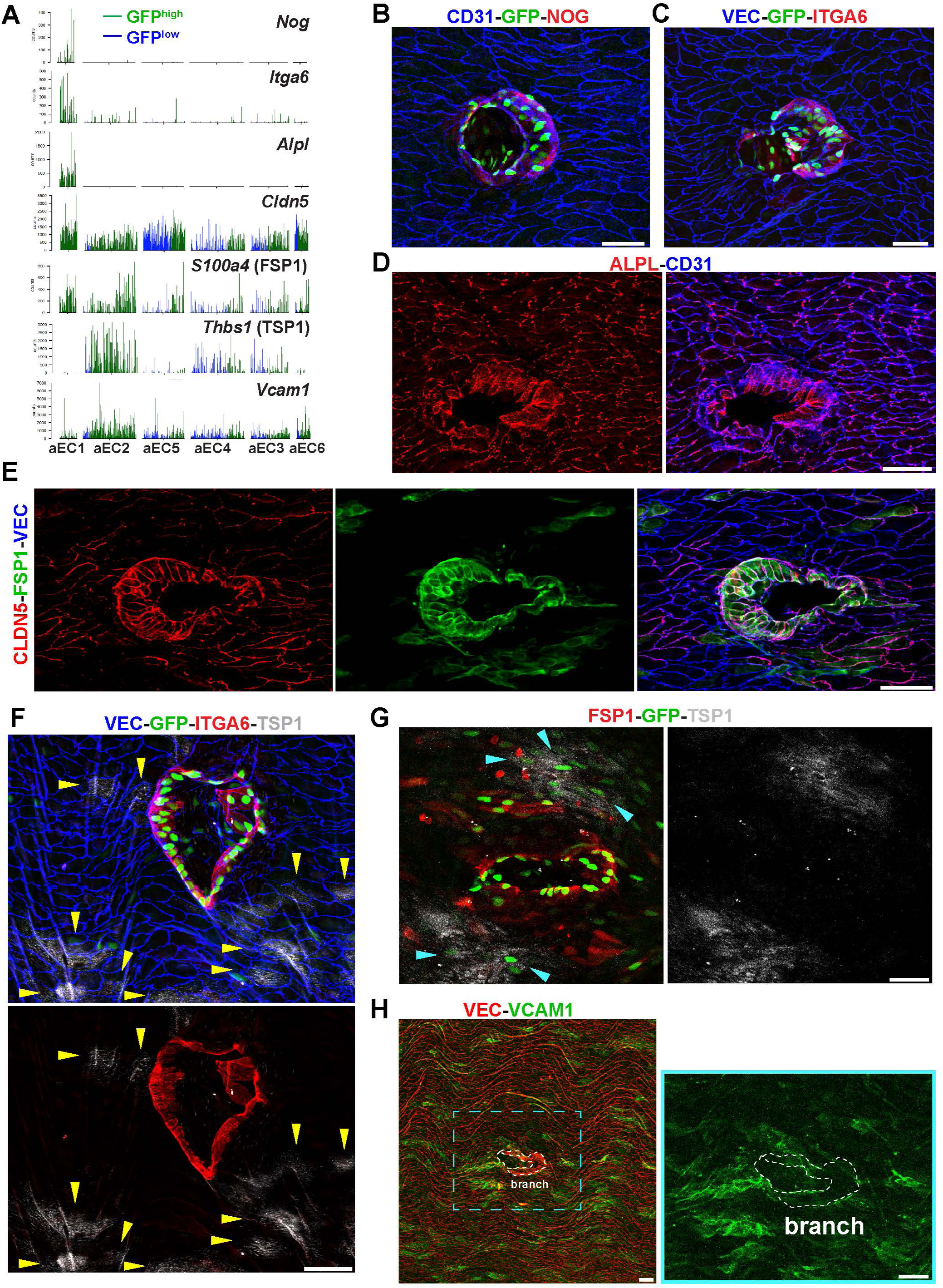
Identification of aEC1 as arterial ECs around the circumference of branch point orifices. (A) Barplots of scRNA-seq read counts for *Nog*, *Itga6*, *Alpl*, *Cldn5*, *S100a4*, *Thbs1*, and *Vcam1* in arterial EC clusters (aEC1-6). Each bar represents a single cell, either GFP^high^ (green bar) or GFP^low^ (blue bar). **(B-H))** Confocal images of whole-mount *en face* preparations of mouse thoracic aortae centered on intercostal branch points. **(B-C)** Immunostaining of S1PR1-GS mouse aortae for Noggin (NOG) and CD31 **(B)**, or Integrin alpha-6 (ITGA6) and VE-Cadherin (VEC) **(C)**. **(D-E)** Immunostaining of C57BL/6J mouse aortae for CD31 and Alkaline phosphatase, tissue- nonspecific isozyme (ALPL) **(D)** or Claudin-5 (CLDN5), S100-A4 (FSP1), and VEC **(E)**. **(F)** Immunostaining of S1PR1-GS mouse aorta for ITGA6, Thrombospondin 1 (TSP1), and VEC. Yellow arrow indicate TSP1+ ECs. **(G)** Immunostaining of an S1PR1-GS mouse aorta for FSP1 and TSP1. The circumference of an intercostal branch point harbors FSP1+GFP+ cells, wereas TSP1+GFP+ cells (cyan arrows) are distal from the circumference of the branch point orifice. **(H)** Immunostaining of a mouse aorta for VCAM1 and VEC. Two rows of cells exhibiting the morphology and VEC-localization of cells around branch point orifices are outlined. Scale bars are 50 µM.

*En face* immunofluorescence staining showed that NOG and ITGA6 were expressed by GFP^high^ MAECs at intercostal branch points (Figure 5B and 5C). These GFP^high^ MAECs comprise the first 2-3 rows of cells around the circumference of branch orifices and include ∼20-30 cells in total. In contrast, cells that are more than 2-3 cells away from branch point orifices did not express NOG or ITGA6 (Figure 5B and 5C). This was also seen for cytoplasmic staining of ALPL (Figure 5D). CLDN5 and FSP1 staining demarcate branch point MAECs but also exhibited heterogeneous staining of surrounding ECs (Figure 5E).

In contrast, TSP1 staining was exluded from ITGA6-expressing GFP^high^ MAECs at branch points (Figure 5F; see also Figure 5-figure supplement 1A). Rather, GFP^high^ MAECs located distally from the circumference of branch points expressed TSP1 in a patchy pattern (Figure 5G). We confirmed the endothelial nature of TSP1 immunoreactivity by staining sagittal sections of thoracic aortae (Figure 5-figure supplement 1B).

Similar to TSP1, VCAM1 staining was low at branch points, while surrounding cells showed heterogeneous levels of immunoreactivity in a somewhat asymmetric manner (Figure 5H). We identified isolated GFP+ cells and patches of GFP+ cells distal from branch points with high VCAM1 immunoreactivity (Figure 5-figure supplement 2A). As expected, intraperitoneal LPS administration induced VCAM1 expression uniformly in aortic ECs (Figure 5-figure supplement 2B). These data suggest that the aEC1 population includes MAECs that are anatomically specific to branch points, while the aEC2 population is located distally and heterogeneously from branch points but nonetheless exhibits S1PR1/ß-arrestin signaling.

### Analysis of branch point-specific arterial ECs cluster aEC1

Having identified the aEC1 cluster as including cells which demarcate thoracic branch points orifices, we sought to characterize these unique cells in further detail. As shown in Figure 4A, aEC1-enriched transcripts are associated with a diverse range of signaling pathways, including MAPK/GPCR, VEGF, and integrin signaling. Among aEC1-enriched transcripts, 16 were up-regulated in *S1pr1*-*ECKO* MAECs, 5 were down-regulated (including *S1pr1*) and the remaining 390 were not differentially expressed (Figure 6A). The 16 *ECKO* up-regulated transcripts included positive regulators of angiogenesis (*Pgf*, *Apold1*, *Itga6*, *Kdr*) (Mirza, Capozzi, Xu, McCullough, & Liu, 2013; Olsson, Dimberg, Kreuger, & Claesson-Welsh, 2006; Primo et al., 2010), regulators of GPCR signaling (*Rgs2*, *Rasgrp3*), and *Cx3cl1* (Fractalkine), which encodes a potent monocyte chemoattractant (White & Greaves, 2012). We noted that several aEC1-enriched transcipts were also expressed in LEC and vEC, but nonetheless were depleted from the remainder of arterial ECs (aEC2-6) (Figure 6B). We examined the 25 transcripts most specific to aEC1 (log_2_ [fold-change] vs. all ECs > 4) and observed that 5 of these (*Dusp26*, *Eps8l2*, *Hapln1*, *Lrmp*, and *Rasd1*) showed differential expression upon loss of S1PR1 function in MAECs (Figure 6C). Therefore, the majority of transcripts specific to aEC1 do not appear to require S1PR1 signaling for normal levels of expression. For example, protein levels of the aEC1 marker ITGA6 were not markedly affected at branch point orifices in *S1pr1-ECKO* animals (Figure 6D). The ∼2-fold increase in *Itga6* transcript levels in *ECKO* MAECs may be due to expression in heterogeneous (non aEC-1) populations, reflecting increases in LECs and/or aEC2-6. Nonetheless, these data suggest that S1PR1 signaling is required for normal expression of some, but not the majority, of transcripts enriched in aEC1 cells located at orifices of intercostal branch points.

**Figure 6.**
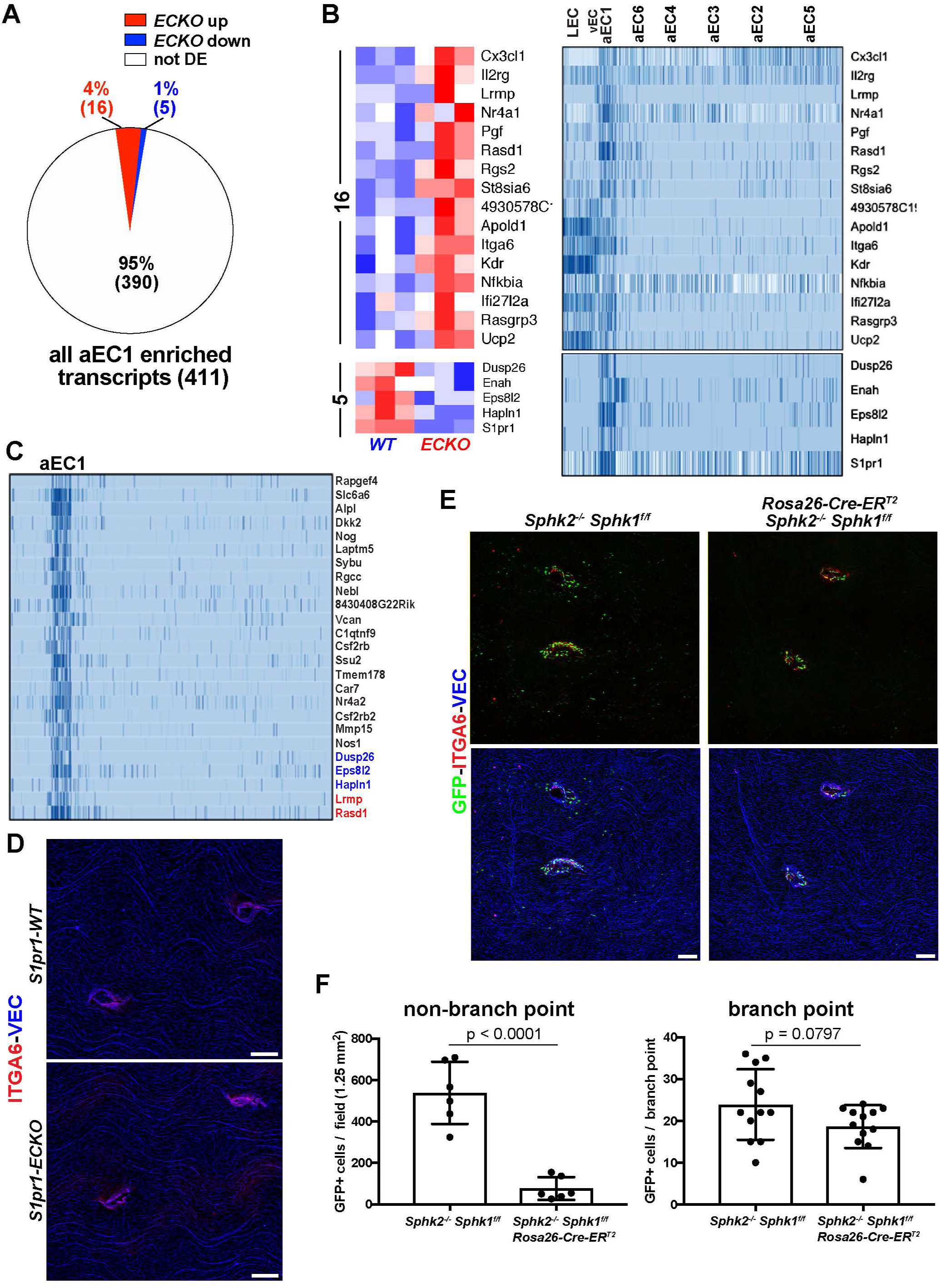
Gene expression in aEC1 cells is largely independent of S1P/S1PR1 signaling. (A) Pie chart of all aEC1-enriched transcripts (versus aEC2-6) indicating those which were also differentially expressed in *S1pr1-ECKO* MAECs. **(B)** Heatmap (row Z-scores) of the 20 transcripts that were both S1PR1-regulated and aEC1-enriched (left). Expression of these transcripts in each cluster from scRNA-seq analysis is shown (right). **(C)** Gene expression in aEC1 was compared to all ECs (LEC, vEC, and aEC2-6 collectively). All transcripts expressed greater than 16-fold higher in aEC1 are shown (25 transcripts total). Red, blue and black transcript names indicate up- regulated, down-regulated or similar levels of expression in *S1pr1-ECKO* MAECs, respectively. **(D)** Immunostaining of *S1pr1-ECKO* and *WT* aortae whole mount *en face* preparations for ITGA6 and VEC. Images are representative of observations from two pairs of animals (N=4). **(E)** ITGA6 and VEC immunostaining of whole mount *en face* preparations of thoracic aortae from S1PR1-GS mice bearing *Sphk2^-/-^ Sphk1*^f/f^ or *Sphk2^-/-^ Sphk1*^f/f^ *Rosa26-Cre-ER^T2^* alleles. **(F)** Quantification of GFP+ arterial EC at branch point (n=12) and non-branch point (n=6) locations (n=2 mice for each genotype). Branch point EC were defined as cells included in the first three rows around the edge of orifices. Only GFP+ EC in the same Z-plane as surrounding arterial EC were counted (GFP+ EC of intercostal arteries were in a different Z-plane and therefore were not counted as branch point EC). All scale bars are 100 µM.

We addressed whether circulatory S1P is required for S1PR1/ß-arrestin coupling in arterial aortic endothelium by genetically deleting the two murine sphingosine kinase enzymes, *Sphk1* and *Sphk2*. We generated S1PR1-GS mice harboring *Sphk1*^f/f^ and *Sphk2*^-/-^ alleles, bred this strain with the tamoxifen-inducible *Rosa26-Cre-ER^T2^* allele, and induced *Sphk1* deletion by tamoxifen injection into adult mice (see Methods). Plasma S1P concentrations in Cre- animals were 631 ± 280 nM, whereas S1P was undetectable in plasma from Cre+ mice. Cre+ mice harbored ∼7-fold fewer non-branch point GFP+ ECs relative to Cre- mice (Figure 6E and 6F). In contrast, the number of branch point GFP+ EC was not significantly different between Cre+ and Cre- animals, suggesting that S1PR1/ß-arrestin coupling (GFP expression) in branch point aEC1 cells is largely independent of S1P-mediated activation of S1PR1 (Figure 6E and 6F). Furthermore, ITGA6 expression at branch point orifices was unaltered in Cre+ mice and therefore is independent of circulatory S1P. Taken together, these data suggest that the unique transcriptome of aEC1 cells is largely independent of S1P/S1PR1 signaling.

For insight into unique regulators of transcription in aEC1 cells, we identified all TFs enriched and depeleted in this cluster (Figure 7-figure supplement 1A; see also Suplementary File 5). Among arterial ECs, TFs highly enriched (Z-score > 7) in aEC1 are *Hey1*, *Nr4a2* (NURR1)*, Nr4a1* (NUR77), *Sox17*, *Ebf1*, and *Bcl6b* (Figure 7A and Figure 7-figure supplement 1A and 1B). *Lef1* transcripts were detected at significant levels only in aEC1 cells (Figure 7-figure supplement 1A and 1B). Transcripts encoding other TFs, such as *Tcf4*, *Ets1*, *Sox18*, *Epas1*, *Mef2c*, and *Tox2*, were aEC1-enriched (Z-score > 3 and < 7) but more heterogeneously distributed in other arterial clusters (Figure 7A and Figure 7-figure supplement 1A). These data are consistent with our ATAC-seq analysis, which showed over-represntation of SOX17, TCF4, and NUR77 motifs in chromatin specifically open in GFP^high^ MAECs. In contrast, *Gata6* (ubiquitous among aEC2-6) and *Gata3* (heterogeneous among aEC2-6) were both depleted from aEC1 (Figure 7A and Figure 7-figure supplement 1A and 1B).

**Figure 7.**
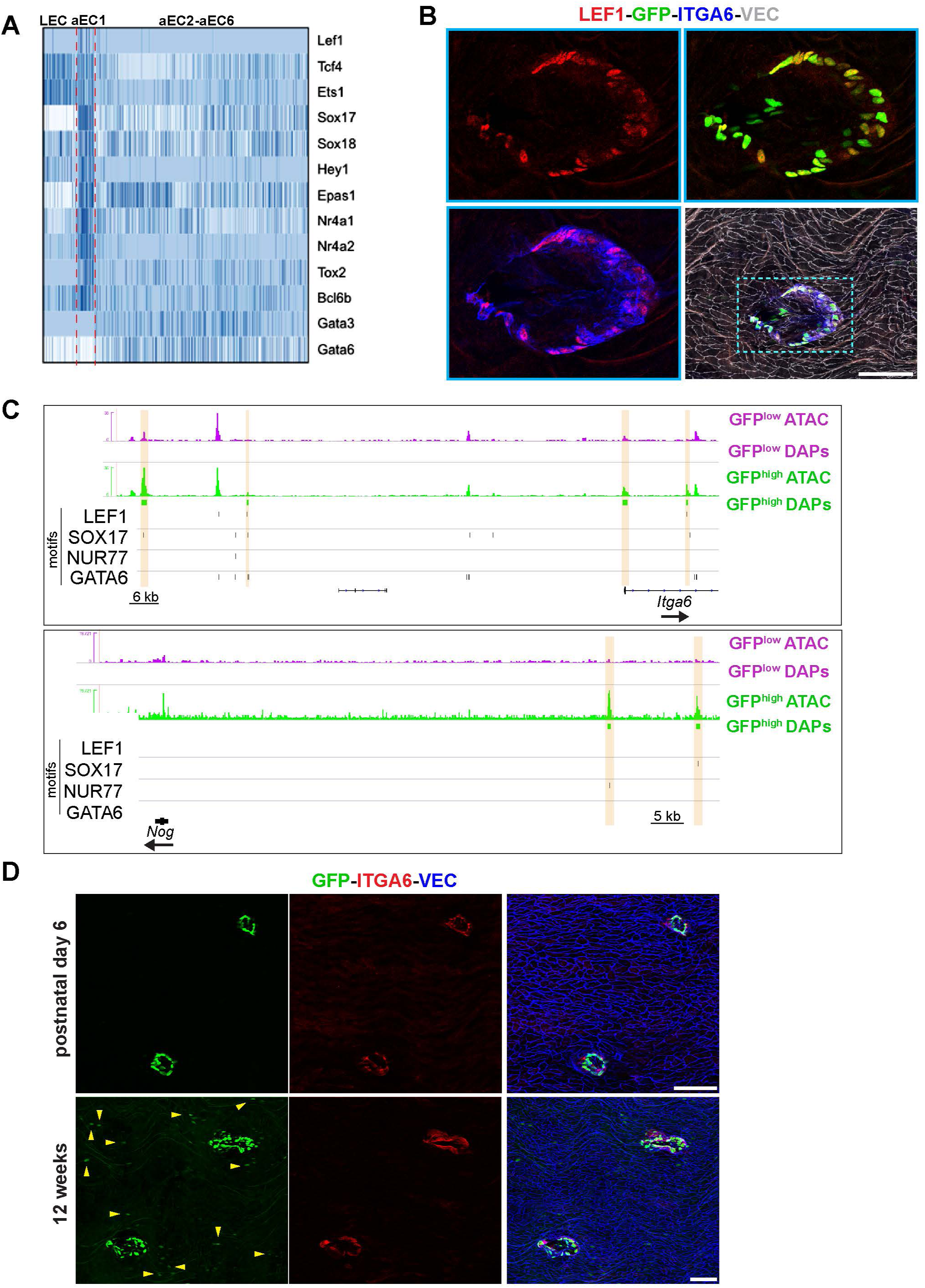
Transcription factors enriched in aEC1 cells. (A) Heatmap of selected transcription factors enriched and depleted in aEC1 cells (see also Figure 7-figure supplement 1A). **(B)** Immunostaining of an S1PR1-GS mouse thoracic aorta for Lymphoid enhancer-binding factor 1 (LEF1), ITGA6, and VEC. Image is representative of observations from 6 total branch points from 2 animals. Scale bar is 100 µM. **(C)** Genome browser images of GFP^high^ and GFP^low^ MAECs ATAC-seq signal (RPGC normalized) at *Itga6* and *Nog* loci. Peaks with increased accessibility in GFP^high^ MAECs (GFP^high^ DAPs) are indicated. All LEF1, Nuclear receptor subfamily 4 group A member 1 (NUR77), Transcription factor SOX-17 (SOX17), and Transcription factor GATA-6 (GATA6) motifs identified in consensus peaks (Figure 2A) are also shown. Orange bars highlight GFP^high^ MAECs DAPs harboring LEF1, SOX17, and/or NUR77 motifs. **(D)** Immunostaining of postnatal day 6 (P6) (n=2) and 12-week (n=4) S1PR1-GS mice thoracic aortae for ITGA6 and VEC. Scale bars are 100 µM.

Immunostaining of thoracic aorta *en face* preparations for LEF1 showed nuclear immunoreactivity in GFP^high^ ECs at branch point orifices but not in adjacent ECs (Figure 7B). We noted that all LEF1+ cells also exhibited ITGA6 expression, confirming these two proteins as markers of aEC1 cells at branch point orifices (Figure 7B). These data suggest involvement of LEF1, a downstream TF of Wnt/ß-catenin signaling, in regulating gene expression in aEC1 cells.

We further explored roles for LEF1, SOX17, NUR77, and GATA6 in aEC1 gene expression by examining binding sites for these factors near genes encoding aEC1-enriched transcripts. A prominent GFP^high^-specific peak in the first intron of *Itga6* harbored a LEF1 motif (Figure 7C). In addition, a GFP^high^-enriched peak ∼115 kb upstream of the transcription start site (TSS) harbored a SOX17 motif. Similarly, we identified bindings sites for SOX17 and NUR77 in GFP^high^-specific peaks upstream of the *Nog* gene (Figure 7C). Each of these putative enhancers lacked GATA6 motifs. In contrast, the *Frzb* and *Pcolce2* genes, which encode aEC4-enriched transcripts and were up-regulated in GFP^low^ MAECs (Figure 1-figure supplement 1B), harbored intronic GFP^low^ MAEC-specific peaks with GATA6 binding sites (Figure 7-figure supplement 2A).

For broader insight into TF activity near aEC1 genes, we extracted all GFP^high^ and GFP^low^ merged peaks that intersected a 100 kb window centered on the TSSs of all aEC1-enriched (Z-score > 3) and -depleted (Z-score < -3) genes. HOMER analysis of these peaks showed enrichment of SOX17, “Ets1-distal”, MEF2C, and NUR77 motifs near the aEC1-enriched genes (Figure 7- figure supplement 1C). In contrast, GATA motifs (GATA2, GATA3, GATA6), as well as the NKX2.2 motif, were enriched near aEC1-depleted genes (Figure 7-figure supplement 1C). Collectively, these data suggest roles for specific transcription factors, such as LEF1, SOX17, NUR77, and GATA6, in mediating transcriptional events which distinguish aEC1 from the remainder of aortic arterial ECs.

Examination of postnatal day 6 (P6) S1PR1-GS aortae showed that MAECs at branch point orifices expressed GFP and ITGA6 in a manner similar to adult S1PR1-GS mice (Figure 7D). We observed that non-branch point GFP+ EC were less frequent in P6 mice, suggesting that heterogeneity among non-branch point ECs changes over time. Indeed, this hypothesis is consistent with McDonald et al. (2019), who reported that aEC3 marker transcripts (e.g. *Atf3*) are absent in aged mice (McDonald et al., 2018). Taken together, these data suggest that aEC2-like (GFP^high^) cells occur at greater frequency as aEC3-like cells disappear over time in the intima of the murine aorta. Furthermore, these P6 data strongly suggest that the GFP^high^ status of MAECs at branch point orifices, as well as their unique gene expression program, is not dynamic throughout postnatal life. Rather, the gene expression specification of these cells likely occurs during development in tandem with epigenetic changes (i.e. chromatin accessibility) and is stable throughout adulthood.

Together, these findings characterize the postnatal, aortic branch point-specific arterial EC subpopulation designated aEC1. These cells have a unique anatomical location and exhibit high S1PR1/ß-arrestin coupling. The transcriptome of this EC subpopulation does not appear to be directly regulated by S1PR1 signaling. Rather, a combination of TFs, such as NUR77, NURR1, SOX17, HEY1, and LEF1, may regulate cluster-defining transcripts in these cells.

### S1PR1 signaling in adventital lymphatic ECs regulates immune and inflammatory gene expression

To locate aorta-associated LEC with high frequency (97%) of S1PR1/ß-arrestin coupling, we utilized antibodies against LYVE1 and VEGFR3 (*Flt4*). LYVE1 marks most but not all LEC subtypes and VEGFR3 is a pan-LEC marker (Wang et al., 2017). Sagittal sections of the S1PR1-GS mouse thoracic aorta revealed that a subset of adventitial LYVE1+ LECs are positive for S1PR1/ß-arrestin coupling (Figure 8A).

**Figure 8.**
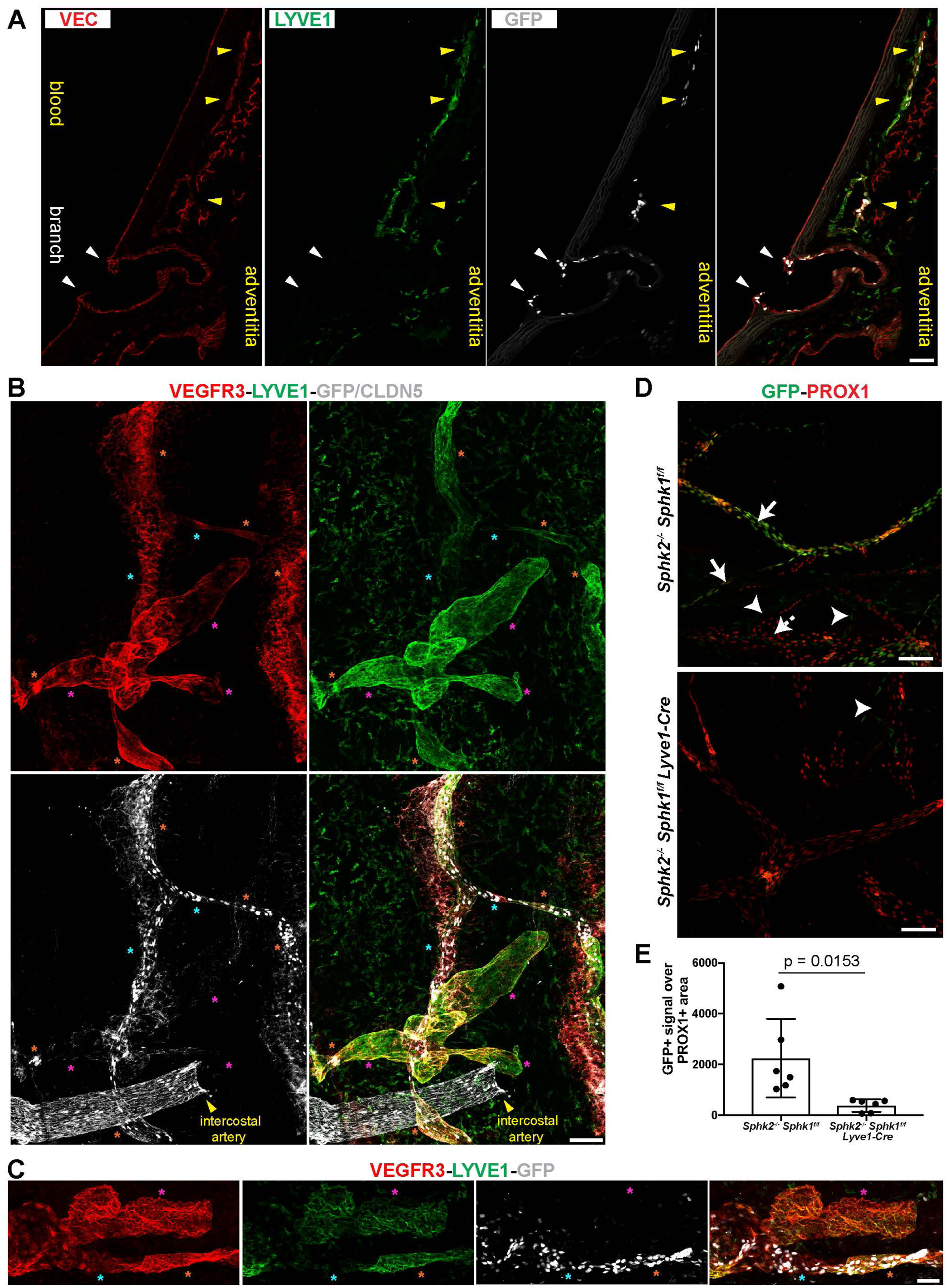
S1P is required for S1PR1/ß-arrestin coupling in LECs of aorta-associated lymphatics. (A) Confocal images of a sagittal cryosection (14 µM) of an S1RP1-GS mouse aorta immunostained for VE-Cadherin and Lymphatic vessel endothelial hyaluronic acid receptor 1 (LYVE1). White arrows indicate GFP+ arterial ECs at a branch point orifices and yellow arrows indicate adventitia-associated GFP+ LECs. Scale bar is 50 µM. **(B-C)** Confocal images of whole- mount preparations of S1RP1-GS mouse thoracic aortae immunostained for Vascular endothelial growth factor receptor 3 (VEGFR3; *Flt4*), LYVE1, and CLDN5 with the tunica intima **(B)** or tunica adventitia **(C)** in contact with coverslip. Orange stars indicate VEGFR3+LYVE1+GFP+ areas, cyan stars indicate VEGFR3+LYVE1^low^GFP+ areas, and magenta stars indicate VEGFR3+LYVE1+GFP- areas. Scale bars are 100 µM **(B)** and 50 µM **(C)**. **(D)** Representative images of mesentery lymphatics from S1PR1-GS mice bearing *Sphk2*^-/-^:*Sphk1*^f/f^ (n=3) or *Sphk2*^-/-^:*Sphk1*^f/f^:*Lyve1-Cre* (n=3) alleles whole mounted and immunostained for PROX1. Solid arrows: GFP+PROX1+, dashed arrows: GFP-PROX1+, arrowheads: GFP+PROX1- (arterioles). Scale bar is 100 µM. **(E)** Quantification of GFP+ signal over PROX1+ areas (N=6). Scale bars are 100 µM.

Next, we prepared whole mounts of thoracic aortae and collected confocal Z-stacks of only the adventitial layer (Figures 8B and 8C). We observed three distinct expression patterns: VEGFR3+LYVE1+GFP+ (orange stars), VEGFR+LYVE1^low^GFP+ (cyan stars), and VEGFR3+LYVE1+GFP- (magenta stars). We noted that the VEGFR3+LYVE1+GFP- areas were associated with blind-ended, bulbous structures. In addition to expression in aEC1, *Itga6* transcripts were detected in vEC and LEC populations (Figure 8-figure supplement 1A). Consistently, we observed GFP+ITGA6+LYVE1+ LEC on the adventitial side of the aorta, in proximity to a GFP+ITGA6+LYVE1- arterial branch point (Figure 8-figure supplement 1B).

We examined the role of LEC-derived S1P in LEC S1PR1/ß-arrestin coupling by generating S1PR1-GS mice deficient in lymph S1P (*S1pr1*^ki/+^:*Sphk1*^f/f^ :*Sphk2*^-/-^:*Lyve1-Cre*^+^) (Pham et al., 2010). LECs were identified in mesenteric vessels by immunostaining for the LEC-specific transcription factor prospero homeobox protein 1 (PROX1). Similar to our observations in the aorta, GFP+PROX1+ cells were heterogeneous, with some PROX1+ structures showing high levels of GFP expression and others showing little to none (Figure 8D). Nonetheless, we observed a significant decrease in the frequency of PROX1+GFP+ LECs in Cre+ mice (Figure 8D and 8E), demonstrating that LEC-derived S1P is required for S1PR1/ß-arrestin coupling in LECs. These data are consistent with abundant LEC expression of the S1P transporter *Spns2* (Figure 3- figure supplement 4A), which is also required for normal levels of lymph S1P (Simmons et al.,2019). Taken together, scRNA-seq analysis of aortic ECs identified two anatomically distinct arterial EC populations (branch point and non-branch point), as well as a subset of adventital LECs, each of which shows high S1PR1/ß-arrestin coupling.

For insight into S1PR1-mediated gene expression in aorta-associated LECs, we divided *S1pr1-ECKO* upregulated genes (Figure 1C and Figure 1-figure supplement 1D) according to their cluster assignments from scRNA-seq analysis. 48% of upregulated genes were enriched in the LEC cluster (Z-score > 3 versus the remainder of ECs), while only 7% were enriched in vEC and/or aEC1-6 (Figure 9A and Suplementary File 6). None of the *S1pr1-ECKO* downregulated transcripts were enriched in the LEC cluster. A heatmap of the 78 LEC transcripts up-regulated in *S1pr1-ECKO* MAECs is shown in Figure 9B. Among these were chemokine/cytokine pathway genes (*Irf8*, *Lbp*, *Il7*, *Il33 Ccl21*, *Tnfaip8l1*) as well as lymphangiogenesis-associated genes (*Kdr, Prox1, Lyve1, Nr2f2*) (Figure 9B and Figure 9-figure supplement 1A). This suggests that loss of S1PR1 signaling in LECs alters transcriptional programs associated with lymphangiogenesis and inflammation/immunity.

**Figure 9.**
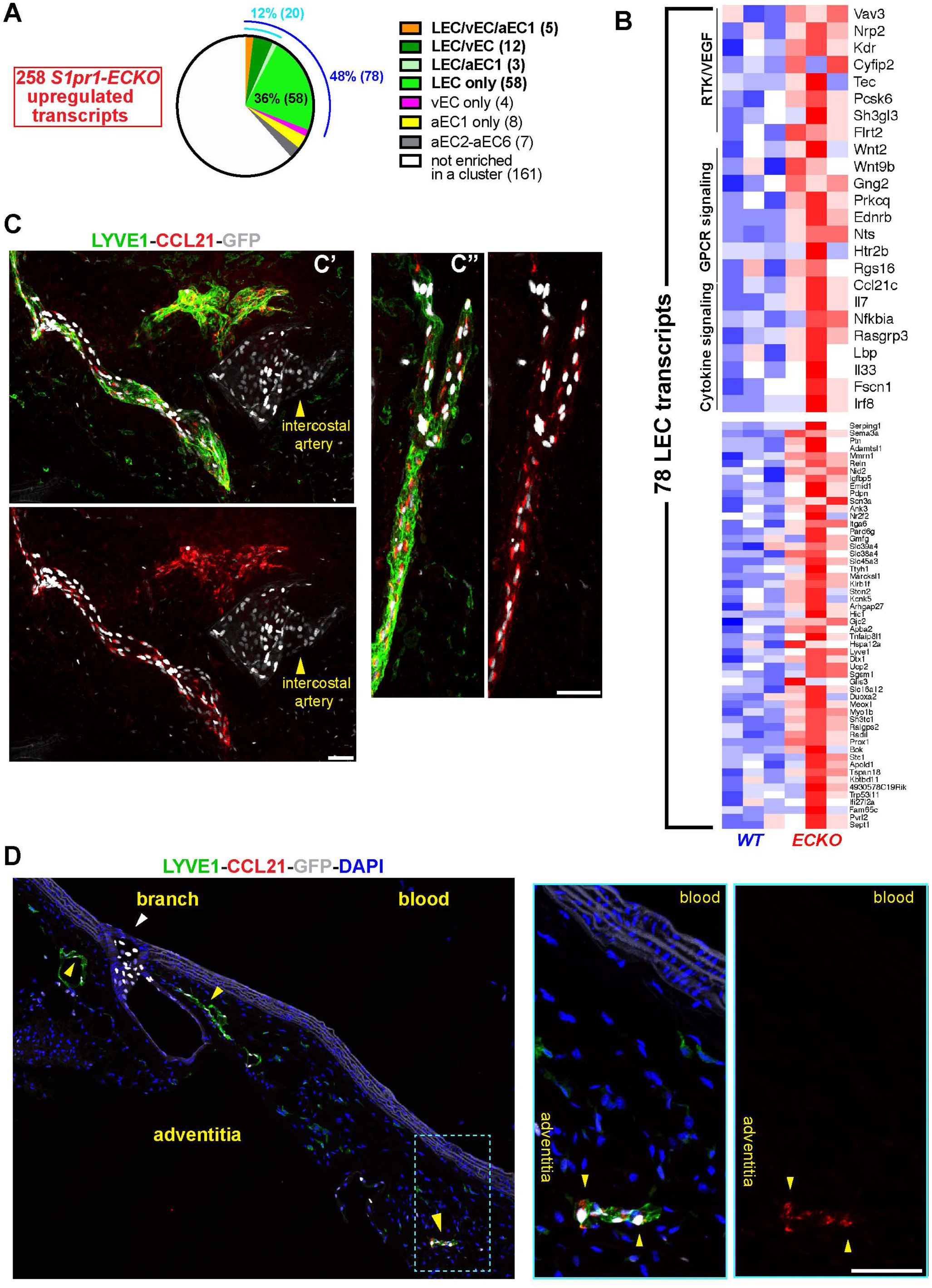
S1PR1 regulation of gene expression in aorta-associated LECs. (**A**) Pie-chart distribution of *S1pr1-ECKO* MAECs up-regulated transcripts. The 258 transcripts were binned according to their cluster assignment from scRNA-seq analysis. The blue and cyan numbers indicate the total percentage of transcripts enriched in LEC and LEC plus at least one cluster, respectively. The numbers in parentheses are absolute transcripts numbers. (**B**) Heatmap (row Z-scores) of all 78 *S1pr1-ECKO* MAECs up-regulated transcripts that are LEC-enriched. Selected GSEA pathways are identified in the top heatmap. (**C**) Confocal images of two fields (c’ and c’’) of a whole-mounted S1RP1-GS mouse thoracic aorta immunostained for LYVE1 and C- C motif chemokine 21 (CCL21). The tunica adventitia was facing the coverslip. **(D)** Confocal images of a sagittal cryosection (14 µM) of an S1RP1-GS mouse thoracic aorta immunostained CCL21 and LYVE1. Yellow arrows indicate adventitia-associated CCL21+LYVE1+GFP+ cells and the white arrow indicate GFP+LYVE1- cells at a branch. Scale bars are 50 µM.

Lymphatic vessels associated with the aorta and large arteries, although not well studied, are thought to be involved in key physiological and pathological processes in vascular and immune systems (Csanyi & Singla, 2019; Galkina & Ley, 2009). For example, LEC expression of CCL21 mediates dendritic cell recruitment to lymphatic vessels during homeostasis and pathological conditions (Vaahtomeri et al., 2017). This chemokine was abundantly expressed in the LEC cluster (Figure 8-figure supplement 1A) and was upregulated by S1PR1 loss-of-function (Figure 9B and Figure 9-figure supplement 1A). Whole mount staining of S1PR1-GS mice thoracic aortae for LYVE1 and CCL21 revealed aorta-associated CCL21+ lymphatics with high levels of S1PR1/ß- arrestin coupling (Figures 9C and 9D). We noted that CCL21 protein appeared as peri-nuclear puncta that likely marks the *trans*-Golgi network, as observed in dermal LECs (Vaahtomeri et al., 2017). Sagittal sections of the thoracic aorta indicate that GFP^high^, LYVE1^+^ adventitial lymphatics express CCL21 protein (Figure 9D), suggesting that ß-arrestin-dependent downregulation of S1PR1 correlates with CCL21 expression (Figure 9D).

These studies show that S1PR1/ß-arrestin coupling in a subset of adventitial lymphatics correlates with S1PR1-mediated attenuation of lymphagiogenic/inflammatory gene expression. We observed that the fraction of PDPN^+^ LECs was not altered between *S1pr1-ECKO* and *WT* aorta tissues (Figure 9–figure supplement 1B), indicating that there is not widespread lymphagiogenesis or LEC proliferation. However, analysis of *Flt4* vs *Pdpn* expression and *Ccl21a* vs *Pdpn* expression showed that there are LECs that express *Flt4* and *Ccl21a* but not *Pdpn* (Figure 9-figure supplement 2). Therefore, it is possible that *S1pr1-ECKO* leads to expansion of a PDPN^low^ population in the adventitia. Nonetheless, these data indicate that S1PR1 mediates gene expression in adventitial lymphatics of the homeostatic adult mouse aorta.

## Discussion

Intracellular signaling through G protein- and ß-arrestin-dependent pathways is tightly regulated at the levels of GPCR expression and ligand availability. Endothelium of major organs, such as brain, lung, skeletal muscle, and the aorta, express unique sets of GPCRs with only 5 receptors commonly expressed, one of which is *S1pr1* (Kaur et al., 2017). Despite ubiquitoius endothelial *S1pr1* expression (Galvani et al., 2015), receptor activation *in vivo*, as reported by GFP in S1PR1-GS mice, revealed heterogeneous signaling in multiple organs (Kono et al., 2014). Here, we describe high frequency of aortic GFP+ (i.e. S1PR1/ß-arrestin coupling) EC around intercostal branch point orifices and heterogeneous GFP+ EC throughout the remainder of intimal aortic endothelium. We and others have shown that endothelial ablation of *S1pr1* (*S1pr1-ECKO*) or reduction of circulatory S1P disrupts endothelial barriers (Camerer et al., 2009; Christensen et al., 2016; Christoffersen et al., 2011; Oo et al., 2011; Yanagida et al., 2017). Furthermore, the descending aorta of *S1pr1-ECKO* mice displayed exacerbated plaque formation in the *Apoe*^-/-^ Western diet (WD)-induced atherosclerosis model (Galvani et al., 2015). However, we lack information regarding EC transcriptional responses that correlate with or are directly downstream of S1PR1 signaling. Here, we profiled the transcriptomes and open chromatin landscapes of aortic ECs with high (GFP^high^) or low (GFP^low^) levels of S1PR1/ß-arrestin coupling, as well as *S1pr1- ECKO* aortic ECs.

*S1pr1-ECKO* MAECs upregulated transcripts in TNFa/cytokine signaling pathways and exhibited enhanced chromatin accessibility at NFkB binding sites. Concomitantly, glucocorticoid receptor pathway was suppressed. These mRNA and chromatin signatures were shared between *S1pr1-ECKO* and GFP^high^ MAECs, suggesting that persistent ß-arrestin recruitment to S1PR1 can result in down-regulation of membrane-localized S1PR1 and a subsequent loss-of-function phenotype. There were many (2,145) DEGs between GFP^high^ and GFP^low^ MAECs, but relatively few (365) between *ECKO* and *WT* MAECs, which suggested that the GFP^high^ and/or GFP^low^ populations reported aortic EC subtypes in addition to S1PR1-regulated transcripts. Indeed, chromatin regions uniquely open in GFP^high^ MAECs were enriched with binding sites for TFs that have well-defined but divergent roles in endothelial cells, such as SOX17 (arterial) (Corada et al., 2013; Zhou, Williams, Smallwood, & Nathans, 2015) and COUP-TFII (venous/lymphatic) (Lindskog et al., 2014). Nonetheless, we present the first collection of putative regulatory regions from freshly isolated mouse aortic ECs, which is a critical dataset for future studies of individual enhancer functionalities.

Our scRNA-seq analysis addressed with high resolution the heterogeneity among MAECs. We identified 6 arterial EC clusters (aEC1-6), 1 lymphatic EC cluster (LEC) and 1 venous EC cluster (vEC). Immunohistochemical analyses revealed LEC cells as including lymphatic structures of the aortic adventita, aEC1 cells as circumscribing intercostal branch point orifices, and aEC2 cells as heterogeneously dispersed throughout intimal endothelium. Each of these clusters harbored high frequency (> 90%) of GFP^high^ EC. We also described aEC3 cells, which contained comparatively few (< 60%) GFP^high^ ECs. aEC3 cells strongly resembled an *Atf3*-positive cluster reported by Mcdonald et al. (2018) that mediates endothelial regeneration (McDonald et al., 2018). Considering that *Atf3*-positive ECs were absent in old (18-month) mice (McDonald et al., 2018), we hypothesize that aEC3-like cells disappear over time while aEC2-like cells increase in frequency in the aorta intima. This notion is supported by the higher frequency of non-branch point GFP^high^ intimal ECs in adult mice relative to P6 pups.

Concordant with findings from two recent aorta scRNA-seq studies (Kalluri et al., 2019; Lukowski et al., 2019), our clustering segregated vEC, LEC, and aEC1 from aEC2-6. We suspect that use of S1PR1-GS mice facilitated deconvolution of LEC and vEC from the distinct aEC1 population. Despite their proximity to intercostal branch points, aEC1 cells do not exhibit a transcriptomic signature prototypical of inflammation, as might be expected (Chiu & Chien, 2011).

Furthermore, high levels of S1PR1/ß-arrestin coupling and expression of unique genes (e.g. *Itga6*) in aEC1 cells are independent of both circulatory S1P and age in postnatal mice. To further explore temporal regulation of cluster-specific genes, we examined scRNA-seq of FACS-sorted VEGFR2^high^ cells from E8.25 embryos (Pijuan-Sala et al., 2019). These embryonic cells exhibited expression of aEC1-enriched transcripts (*Lef1*, *Itga6*, *Rasgrp3*, *Alpl*, *Hey1*, *Flt1*, *Igf2*), but depletion of aEC2-6-enriched transcripts (*Thbs1*, *Sod3*, *Vcam1*, *Sfrp1*, *Ace*, *Bmp6*, *Dcn*). This suggests that aEC1 cells are more characteristic of embryonic EC than are the majority of intimal ECs. Similarly, vEC/LEC-specific transcripts (*Prox1*, *Nr2f2*, *Kdr*) were also expressed in these embryonic cells. Thus, transcriptomic similarities between aEC1 and LEC/vEC in the adult aorta may be retained from development, perhaps through epigenetic modifications common between these cell types. It is also possible that the unique anatomical location of aEC1 (at the circumferential ridge at aortic branch point) may promote a distinct EC phenotype either because of spatial positioning or environmental factors.

*S1pr1-ECKO* and *S1pr1*^-/-^ animals display an aortic hyper-branching phenotype between E11.5 and E13.5 that is incompatible with life after E14.5 (Gaengel et al., 2012). Therefore, S1PR1 is required for normal embryonic branching morphogenesis. Consistently, E9.5-E10.5 S1PR1-GS embryos show high GFP expression (S1PR1/ß-arrestin coupling) throughout the dorsal aorta (Kono et al., 2014). This contrasts with the adult aortae, wherein the highest levels of S1PR1/ß-arrestin coupling are concentrated around the orifices of intercostal branch points. These findings further suggest that that the unique gene expression program of aEC1 cells is established during morphogenesis of intercostal arteries during development.

The extent to which aEC1 cells at branch point orifices are functionally distinct remains to be determined. Identification of ITGA6 as a marker of this population will facilitate future studies. For example, combinations of pan-EC, LEC, and ITGA6 antibodies can be used to purify or enrich this population in developmental or disease models (e.g. atherosclerosis). Moreover, *cis*-elements proximal to aEC1-specific genes can be applied to the “Dre-rox/Cre-loxP” system (Pu et al., 2018) to specifically manipulate gene expression in aEC1 cells.

Non-branch point (i.e. aEC2) cells required circulatory S1P for normal levels of S1PR1/ß- arrestin coupling. Similarly, mesenteric LECs required S1P in lymph for normal levels of S1pr1/ß- arrestin coupling. Notably, we did not detect *S1pr1-ECKO* down-regulated transcripts in aEC2 or LEC clusters, but we did detect 4 such transcripts in aEC1 cells, which were *Dusp26*, *Enah*, *Eps8l2*, and *Hapln1*. Among the clusters identified in this study, LEC-enriched transcripts were affected the most upon EC ablation of *S1pr1*. This suggests that loss of S1P/S1PR1 signaling either alters cell-intrisic phenotypes of peri-aortic LECs or induces expansion of one or multiple LEC subtypes.

There is accumulating direct and indirect evidence for key roles of adventital lymphatics in atherogenesis (Csanyi & Singla, 2019; Maiellaro & Taylor, 2007). For example, auto-antibodies against oxidized LDL (OxLDL) inhibit macrophage OxLDL uptake and mitigate atherosclerosis (Shaw et al., 2000). This implies that antigen presenting cells (APCs) phagocytose OxLDL epitopes, then travel via adventitial lymphatics to lymphoid organs (e.g. lymph nodes) and present OxLDL antigens to B cells. Murine atherosclerotic lesions were found to harbor “atypical, lymphatic-like” capillaries that were VEGFR3+ but LYVE1- (Taher et al., 2016), which is consistent with our observations of adventitial LEC heterogeneity. Considering the critical role of lymphatic EC-derived CCL21 in regulating the trafficking of APCs (Vaahtomeri et al., 2017), and perhaps other adaptive immune cells, there is an impetus to determine the extent to which advential lymphatics are a viable target for atherosclerosis therapy.

While there is scant information about the roles of S1P/S1PR1 signaling in adult lymphatic vasculature, our findings lay a groundwork for future studies of S1PR1-mediated LEC phenotype regulation in homeostatic processes and inflammatatory/ autoimmune diseases. A recent study found that lymph-derived S1P facilitates CCL21 deposition in high endothelial venules and dendritic cell recruitment (Simmons et al., 2019). While *S1pr1-ECKO* animals exhibit exacerbated atherosclerosis (Galvani et al., 2015), we cannot discern whether this was due to phenotypes of lymphatic ECs, arterial ECs, or both cell types. Future studies should use artery- and lymphatic-specific Cre-drivers to distinguish between the roles of S1PR1 in different types of vasculature. Such mechanistic studies will help to determine the utility of S1PR1 modulators in treating lymphatic-mediated vasculopathies.

## Methods

### Mice

Animal experiment protocols were approved by the Institutional Animal Care and Use Committees (IACUC) of Boston Children’s Hospital and the French Department of Eduction. S1PR1-GS mice were previously reported (Kono et al., 2014). S1PR1-GS mice used for experiments harbored a single *S1pr1^knockin^* (*S1pr1-tTA-IRES-mArrb2-TEV*) allele (*S1pr1*^ki/+^) as well as a single *H2B-GFP* reporter allele. *S1pr1*^f/f^ mice (Allende, Yamashita, & Proia, 2003) were bred with *Cdh5*-*Cre^ERT2^* mice (Sorensen, Adams, & Gossler, 2009) to generate *S1pr1-ECKO* mice, as previously described (Galvani et al., 2015) (Jung et al., 2012). Gene deletion was achieved by intraperitoneal injection of tamoxifen (2 mg/day) at 5-6 weeks of age for five consecutive days. Tamoxifen treated mice were rested for a minimum of 2 weeks prior to experiments.

S1PR1-GS mice deficient in LEC S1P production were generated by excising a conditional knockout allele for *Sphk1* in an *Sphk2* knockout background with *Lyve1-Cre* (*S1pr1*^ki/+^:*Sphk1*^f/f^:*Sphk2*^-/-^:*Lyve1-Cre*^+^) essentially as described by Cyster and colleagues (Pham et al., 2010) and excision efficiency confirmed by the induction of lymphopenia. S1PR1-GS mice deficient in plasma S1P (S1PR1-GS-S1P-less) were generated by crossing S1PR1-GS mice with *Sphk1*^f/f^:*Sphk2*^-/-^:*Rosa26-Cre-ER^T2^* mice to obtain *S1pr1*^ki/+^:*GFP*^+^:*Sphk1*^f/f^:*Sphk2*^-/-^:*Rosa26-Cre- ER^T2^*^+^ mice (the *Rosa26-Cre-ER^T2^* allele is described in (Takeda, Cowan, & Fong, 2007)). Tamoxifen was administered to S1PR1-GS-S1P-less mice and Cre- littermate controls as described above. Experiments were performed between 23 and 25 weeks after the final tamoxifen dose.

Adult (aged 8 to 12 weeks) males and females were used for sequencing experiments. Males and females of similar age (7 to 18 weeks) were used for imaging studies, unless indicated otherwise. For examination of VCAM1 (Figure 5-figure supplement 2B), 200 *µ*L lipopolysaccharide (Sigma, L2630) in PBS was injected i.p. (5.5 mg/kg) for nine hours followed by euthanasia and tissue harvest.

### FACS isolation and single cell sequencing of mouse aortic endothelial cells

After CO_2_ euthanasia, the right atrium was opened and the left ventricle was perfused with 10 mL phosphate-buffered saline (PBS) (Corning). Aortae were dissected from the root to below the common iliac bifurcation and transferred into ice ice-cold 1x HBSS (Sigma, H1641). Whole aortae were incubated in HBSS containing elastase (4.6 U/mL, LS002292, Worthington), dispase II (1.3 U/mL, Roche), and hyaluronidase (50.5 U/mL, Sigma, H3506) at 37°C for 10 minutes in wells of a 6-well plate. Aortae were then transferred to a 100 mm dish with 1 mL HBSS and minced using small scissors. Minced aortae were transferred to a low protein binding 5 mL tube containing Liberase (0.6 U/mL, Sigma), collagenase II (86.7 U/mL, LS004174, Worthington), and DNase (62.0 U/mL, Sigma, D4527) in 4.3 mL HBSS and incubated at 37°C for 40 minutes with rotation in a hybridization oven. The cell suspension was then triturated 10 times through an 18 G needle to dissociate clumps, followed by addition of 400 *µ*L STOP solution (3 mM EDTA, 0.5% fatty acid-free bovin serum albumin (FAF-BSA) (Sigma, A6003) in 1x HBSS). For the remainder of the procedure, cells were kept on ice and all centrifugation steps were performed at 4°C.

Cells were spun at 500 xg for 5 minutes, the supernatant was removed, then cells were washed with 4 mL STOP solution and spun at 500 xg for 5 minutes. The supernatant was removed, then cells were washed with 4 mL blocking solution (0.25% FAF-BSA in HBSS) and filtered through FACS tubes with filter caps (Falcon). After centrifugation and supernatant removal, cells were stained with phycoerythrin (PE)-conjugated anti-mouse CD31 (MEC13.3, Biolegend, San Diego, CA), allophycocyanin (APC)-conjugated anti-mouse CD45 (30-F11, Biolegend) and APC- conjugated TER119 (Biolegend, 116212) antibodies in blocking solution with anti-CD16/32 (2.5 *µ*g/mL) for 45 minutes on ice. DAPI (0.7 *µ*M) was added for the final 5 minutes of staining to exclude dead cells. Aortic cells were washed with 4.5 mL FACS buffer (0.25% FAF-BSA in PBS) before sorting for selection of CD31^+^/CD45^-^/TER119^-^/GFP^high^ and CD31^+^/CD45^-^/TER119^-^

/GFP^low^ cells using BD FACSAria^TM^ II (BD Bioscience) (see Figure 1B). Cells from *S1pr1*-WT and -*ECKO* mice were sorted using the GFP^low^ gate (see Figure 1-figure supplement 1A) because it includes MAECs from mice genetically negative for the *H2B-GFP* reporter allele and stained with the same antibody panel. Cells from 2-4 aortae of age and sex-matched adult mice were pooled for each individual experiment (ATAC-seq, RNA-seq and scRNA-seq). Cells were sorted into either 0.1% FAF-BSA/PBS or buffer RLT (Qiagen) supplemented with b-mercaptoethanol for ATAC-seq and RNA-seq, respectively.

For single-cell RNA-seq, GFP^high^ and GFP^low^ cells were gated as described above. Library preparation from single cells was performed as previously described (Vanlandewijck et al., 2018). Briefly, cells were deposited into individual wells of 384-well plates containing 2.3 *µ*L of lysis buffer (0.2% Triton-X (Sigma, T9284), 2U/µL RNase inhibitor (ClonTech, 2313B), 2 mM dNTP’s (ThermoFisher Scientific, R1122), 1 µM Smart-dT30VN (Sigma), ERCC 1:4 × 10^7^ dilution (Ambion, # 4456740)) prior to library preparation using the Smart-seq2 protocol (Picelli et al., 2014).

### Bulk RNA-seq and analysis

Cells sorted into buffer RLT were subjected to total RNA extraction using the RNeasy Micro Kit (Qiagen). The High Sensitivity RNA ScreenTape (Agilent) was used to verify RNA quality before synthesis of double-stranded cDNA from 5-10 ng RNA using the SMART-Seq2 v4 Ultra Low RNA Kit for Sequencing (Takara) according to the manufacturer’s instructions. Agilent 2100 Bioanalyzer and High Sensitivity DNA Kit (Agilent) were used to verify cDNA quality. cDNA libraries were prepared for sequencing using the Illumina Nextera XT2 kit (Illumina), and ∼20-40 million paired-end reads (2 x 75 bp) were sequenced for each sample.

Reads from each sample were aligned to the MGSCv37 (mm9) genome assembly using STAR (Dobin et al., 2013) with the options: --runMode alignReads --outFilterType BySJout --outFilterMultimapNmax 20 --alignSJoverhangMin 8 --alignSJDBoverhangMin 1 --outFilterMismatchNmax 999 --alignIntronMin 10 --alignIntronMax 1000000 -- alignMatesGapMax 1000000 --outSAMtype BAM SortedByCoordinate --quantMode TranscriptomeSAM. Gene-level counts over UCSC annotated exons were calculated using the Rsubread package and “featureCounts” script (Liao, Smyth, & Shi, 2013) with options: -M –O –p –d 30 –D 50000. The resultant count table was input to edgeR (M. D. Robinson, McCarthy, & Smyth, 2010) for differential gene expression analysis. The .bam files from STAR were input to the RSEM (B. Li & Dewey, 2011) script “rsem-calculate-expression” with default parameters to generate FPKMs for each replicate.

### ATAC-seq analysis

ATAC-seq libraries were prepared according to the previously described fast-ATAC protocol (Corces et al., 2016). Briefly, 800-4,000 FACS-isolated cells in 0.1% FAF-BSA/PBS were pelleted by centrifugation at 400 × *g* at 4°C for 5 min. Supernatant was carefully removed to leave the cell pellet undisturbed, then cells were washed once with 1 mL ice-cold PBS. The transposition mix [25 µL buffer TD, 2.5 µL TDE1 (both from Illumina FC-121-1030), 1 µL of 0.5% digitonin (Promega, G9441) and 16 µl nuclease-free water] was prepared and mixed by pipetting, then added to the cell pellet. Pellets were disrupted by gently flicking the tubes, followed by incubation at 37°C for 30 minutes in an Eppendorf ThermoMixer with constant agitation at 300 rpm. Tagmented DNA was purified using the MinElute Reaction Cleanup Kit (Qiagen, 28204), and subjected to cycle-limiting PCR as previously described (Buenrostro et al., 2013). Transposed fragments were purified using the MinElute PCR Purification Kit (Qiagen, 28004) and Agilent DNA Tapestation D1000 High Sensitivity chips (Agilent) were used to quantify libraries. ∼20-60 million paired-end reads (2 x 75 bp) were sequenced for each sample on a NextSeq instrument (Illumina).

Read alignment to the MGSCv37 (mm9) genome assembly was performed with bowtie2 (Langmead & Salzberg, 2012) and the options: --very-sensitive –X 2000 –no-mixed –no-discordant. Duplicated fragments were removed using the Picard “MarkDuplicates” script with the options: Remove_Duplicates=true Validation_stringency=lenient (http://broadinstitute.github.io/picard/).

Paired-end reads were separated, centered on Tn5 cut sites, and trimmed to 10 bp using a custom in-house script. Peaks were called using the MACS2 “callpeak” script (Zhang et al., 2008) with options: -B –keep-dup all –nomodel –nolambda –shift -75 –extsize 150. Reads mapping to murine blacklisted regions and mitochondrial DNA were masked out of peak lists using the Bedtools “intersect -v” script (Quinlan & Hall, 2010).

Replicates from each biological group were merged using the bedops “merge” script to generate one high-confidence peak set for each of the four biological groups (GFP^high^, GFP^low^, *S1pr1*-*ECKO*, *S1pr1*-*WT*) (Neph et al., 2012). These four peak sets were then merged to generate a merged, consensus peak set of 123,473 peaks. For each replicate, reads covering consensus peak intervals were counted using the Bedtools “coverage” script with the “-counts” option (Quinlan & Hall, 2010). The resultant count table was input to edgeR (M. D. Robinson et al., 2010) to determine differentially accessible peaks (DAPs).

DAPs were used as input for the HOMER “findMotifsGenome.pl” script with the option “-size given” to identify motifs enriched in peaks with enhanced accessibility in either GFP^high^, GFP^low^, *S1pr1-ECKO*, or *S1pr1-WT* MAECs (Heinz et al., 2010).

Nucleotide-resolution coverage (bigwig) tracks were generated by first combining trimmed reads from each replicate, then inputting the resultant .bam files to the DeepTools (Ramirez et al., 2016) “bamCoverage” script with options “—normalizeUsing RPGC –binSize 1”. Heatmaps of ATAC-seq reads within 600 bp of p65, NUR77, COUP-TFII, ATF1, GATA2, and GRE motifs were generated by centering coverage tracks on each motif identified in DAPs. These motifs were identified using the HOMER script “annotatePeaks.pl” with the “-m -mbed” options. All heatmaps were generated using DeepTools and all genome browser images were captured using Integrative Genomics Viewer (J. T. Robinson et al., 2011).

### scRNA-seq analysis

1152 Fastq files (one per cell) were aligned to the GRCm38 (mm10) genome assembly using STAR with options –runThreadN 4 –outSAMstrandField intronMotif –twopassmode Basic. Bam files were input to the velocyto (http://velocyto.org/velocyto.py/) command-line script “run-smartseq2” (La Manno et al., 2018). Expressed repetitive elments were downloaded from the UCSC genome browser and masked from analysis using the “-m” option of the “run-smartseq2” script. The resultant table of read counts per transcript (“loom” file) was input to the PAGODA2 (https://github.com/hms-dbmi/pagoda2) R package for further analysis (Fan et al., 2016; La Manno et al., 2018). The details of our R code are provided in Supplementary File 9.

After variance normalization, the top 3000 overdispersed genes were used for principal component analysis (PCA). An approximate k-nearest neighbor graph (k = 30) based on a costine distance of the top 100 principal components was used for clustering. Cluster were determined using the multilevel community detection algorithm. PCA results were plotted using the “tSNE” embedding option of the PAGODA2 “r$getEmbedding” function. Heatmaps of gene expression embedded on hierarchical clustering, differential expression analyses, and expression of individual transcripts on the tSNE embedding were generated using the graphical user interface at http://pklab.med.harvard.edu/nikolas/pagoda2/frontend/current/pagodaLocal/. We generated the binary (.bin) file according to the Pagoda2 walkthrough: https://github.com/hms-dbmi/pagoda2/blob/master/vignettes/pagoda2.walkthrough.oct2018.md. This binary file (Supplementary File 7) can be uploaded to the graphical user interface for exploration of our dataset. We generated a file of cluster labels (for LEC, vEC, VSMC, aEC1, aEC2, aEC3, aEC4, aEC5, aEC6 as well as all other cluster grouping used for analysis), which can also be uploaded to the graphical user interface for visualization of these clusters (Supplementary File 8).

### Immunohistochemistry

Mice were euthanized as described above, then perfused through the left ventricle with 5 mL PBS immediately followed by 10 mL ice-fold 4% paraformaldehyde (PFA) in PBS. The left ventricle was then perfused with 6 mL PBS. After aorta dissection, remaining fat tissue was removed with the aorta suspended in PBS in a polystyrene dish. For sectioning, intact thoracic aortae were additionally post-fixed in 4% PFA for 10 minutes, then briefly washed with PBS three times. Aortae were then cryoprotected in 30% sucrose in PBS for 2 hours at 4° C, embedded in a 1:1 mixture of 30% sucrose PBS:OCT over dry ice, then sectioned at 14 *µ*M intervals using a cryostat (Leica Biosystems).

For *en face* preparations, fine scissors were used to cut the aorta open and expose the endothelium for downstream flat-mount preparation. Aortae were placed into 24-well tissue culture plates and permeabilized in 0.5% Triton X-100/PBS (PBS-T) for 30 minutes on a room-temperature orbital shaker, then blocked in blocking solution (1% BSA (Fisher Scientific, BP1605), 0.5% Donkey Serum (Sigma, D9663) in PBS-T for 1 hour. Primary antibody incubations were carried out overnight in blocking solution, followed by detection with secondary antibodies. Primary antibodies used were goat anti-VE-cadherin (1:300, R&D systems, AF1002), rabbit anti- LYVE1 (1:300, 103-PA50AG, ReliaTech), goat anti-LYVE1 (1:300, R&D systems, AF2125), goat anti-VEGFR3 (1:200, R&D systems), rat anti-ITGA6 (1:200, BioLegend, 313602) goat anti-ALPL (1:200, R&D systems, AF2910), biotinylated mouse-anti TSP1 (1:200, ThermoFisher, MA5-13395), Alexa Fluor 488-conjugated mouse anti-CLDN5 (1:100, Invitrogen, 352588), rabbit anti-FSP1 (1:300, MilliporeSigma, 07-2274), goat anti-CD31 (1:300, R&D systems, AF3628), rat anti-CD31 (1:100, HistoBiotec, DIA-310), rabbit anti-LEF1 (1:200, Cell Signaling Technologies, 2230), rat anti-VCAM1 (1:200, MilliporeSigma, CBL1300), goat anti-NOG (1:200, R&D systems, AF719), goat anti-CCL21 (1:200, R&D systems, AF457), goat anti-PROX1 (1:200, R&D systems, AF2727). Following primary antibody incubation, aortae were washed three times in PBS-T for 20 minutes each, then incubated with secondary antibodies in blocking solution at room temperature for 90 minutes. Donkey anti-rat, anti-rabbit, and anti-goat secondary antibodies were purchased from ThermoFisher or Jackson ImmunoResearch as conjugated to Alexa Fluor 405, 488, 546, 568, 594, or 647. TSP1 was detected with streptavidin from Jackson Immunoresearch conjugated to Alexa Fluor 594 or 647. After secondary antibody incubation, aortae were washed in PBS-T for 20 minutes four times, then once in PBS, then mounted in mounting reagent (ProLong® Gold, Invitrogen) on a slide with the tunica intima in contact with the coverslip. A textbook-sized weight was placed on top of the coverslip for 1 minute before sealing with nail polish. Mesenteric vessels were whole-mounted and stained as described above.

### Confocal microscopy and image analysis

Images were acquired using a Zeiss LSM810 confocal microscope equipped with an Plan- Apochromat 20x/0.8 or a Plan-Apochromat 40x/1.4 oil DIC objective. Images were captured using Zen2.1 (Zeiss) software and processed with Fiji (NIH). Zen2.1 software was used to threshold GFP signal and manually count GFP+ nuclei per field (Figure 6). Fiji was used to quantify GFP signal over PROX1+ areas (Figure 8). Figures were assembled in Adobe Illustrator.

### S1P sample preparations

Plasma S1P was extracted as previously decribed (Frej et al., 2015) with minor modification. Plasma aliquots (5 or 10 *µ*L) were first diluted to 100 *µ*L with TBS Buffer (50 mM Tris-HCl pH 7.5, 0.15 M NaCl). S1P was extracted by adding 100 *µ*L precipitation solution (20 nM D7-S1P in methanol) followed by 30 seconds of vortexing. Precipitated samples were centrifuged at 18,000 rpm for 5 minutes and supernatant were transferred to vials for LC-MS/MS analysis (see below).

C18-S1P (Avanti Lipids) was dissolved in methanol to obtain a 1 mM stock solution. Standard samples were prepared by diluting the stock in 4% fatty acid free BSA (Sigma) in TBS to obtain 1 *µ*M and stored at -80 °C. Before analysis, the 1 *µ*M S1P solution was diluted with 4% BSA in TBS to obtain the following concentrations: 0.5 *µ*M, 0.25 *µ*M, 0.125 *µ*M, 0.0625 *µ*M, 0.03125 *µ*M, 0.0156 *µ*M, and 0.0078 *µ*M. S1P in diluted samples (100 *µ*L) were extracted with 100 *µ*L of methanol containing 20 nM of D7-S1P followed by 30 seconds of vortexing. Precipitated samples were centrifuged at 18,000 rpm for 5 minutes and the supernatants were transferred to vials for LC-MS/MS analysis. The internal deuterium-labeled standard (D7-S1P, Avanti Lipids) was dissolved in methanol to obtain a 200 nM stock solution and stored at -20 °C. Before analysis, the stock solution was diluted to 20 nM for sample precipitation.

### LC-MS/MS S1P measurement and data analysis

The samples were analyzed with Q Exactive mass spectrometer coupled to a Vanquish UHPLC System (Thermo Fisher Scientific). Analytes were separated using a reverse phase column maintained at 60 °C (XSelect CSH C18 XP column 2.5 *µ*m, 2.1 mm X 50 mm, Waters). The gradient solvents were as follows: Solvent A (water/methanol/formic acid 97/2/1 (v/v/v)) and

Solvent B (methanol/acetone/water/formic acid 68/29/2/1 (v/v/v/v)). The analytical gradient was run at 0.4 mL/min from 50-100% Solvent B for 5.4 minutes, 100% for 5.5 minutes, followed by one minute of 50% Solvent B. A targeted MS2 strategy (also known as parallel reaction monitoring, PRM) was performed to isolate S1P (380.26 m/z) and D7-S1P (387.30 m/z) using a 1.6 m/z window, and the HCD-activated (stepped CE 25, 30, 50%) MS2 ions were scanned in the Orbitrap at 70 K. The area under the curve (AUC) of MS2 ions (S1P, 264.2686 m/z; D7-S1P, 271.3125 m/z) was calculated using Skyline (MacLean et al., 2010).

Quantitative linearity was determined by plotting the AUC of the standard samples (C18- S1P) normalized by the AUC of internal standard (D7-S1P); (y) versus the spiked concentration of S1P (x). Correlation coefficient (R2) was calculated as the value of the joint variation between x and y. Linear regression equation was used to determined analyte concentrations.

## Acknowledgements

We thank Boston Children’s Hospital & Harvard Stem Cell Institute Flow Cytometry Research Facility and Harvard Medical School Biopolymers Facility for technical assistance. We also thank members of the Hla laboratory for critical comments on the project and Dr. Sylvain Galvani for technical advice regarding mouse aorta dissection. This work is supported by NIH grants (R35- HL135821 to T.H.), Fondation Leducq Transatlantic Network grant (SphingoNet) to T.H., C.B., E.C. and R.L.P., intramural research program of NIDDK intramural program grants (R.L.P) and a postdoctoral fellowship from the American Heart Association to A.K.

---

**Figure 1 supplement 1.**
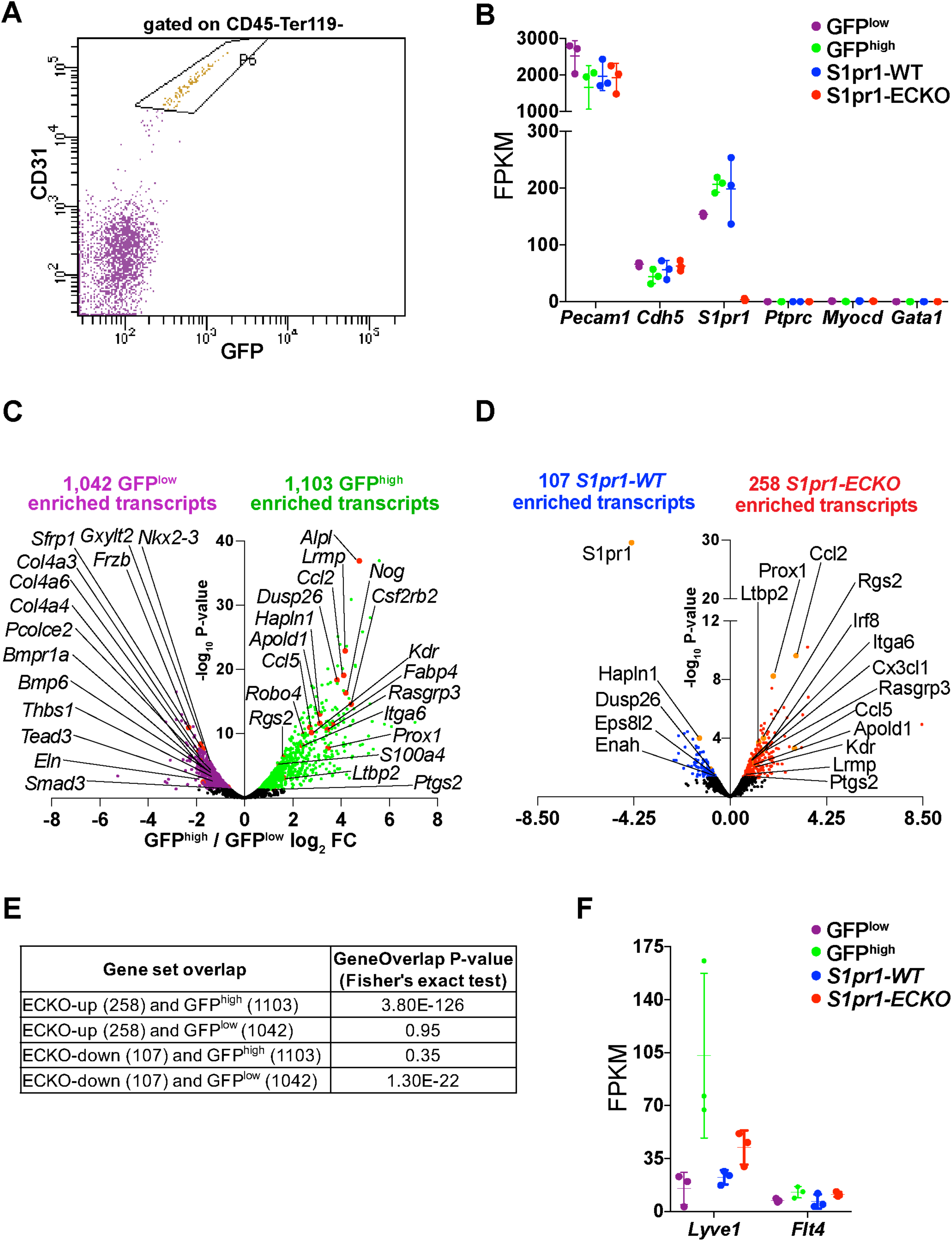
RNA-seq quality control and differential gene expression between GFP^high^ and GFP^low^ and *S1pr1-ECKO* and *WT* MAECs. (A) FACS gating scheme for isolation of *S1pr1-WT* and *S1pr1-ECKO*. MAECs were defined as cells in the gate labeled “P6”, (CD45^-^Ter119^-^CD31^+^). **(B)** Expression, in fragments per kilobase per million mapped fragments (FPKMs), of indicated transcripts from bulk RNA-seq of GFP^high^, GFP^low^, *S1pr1-ECKO* and *WT* MAECs. **(C-D)** Volcano plots of all transcripts (counts per million > 1) illustrate those differentially expressed between GFP^high^ and GFP^low^ **(C)** and *S1pr1-ECKO* and *WT* (D) MAECs (see Supplementary file 1). Highlighted transcripts are associated with TGFß signaling, inflammatory pathways, and/or show common regulation in GFP^high^ and *S1pr1-ECKO* MAECs. **(E)** GeneOverlap (Shen et al., 2013) results for the overlaps shown in Figure 1C. **(F)** Expression of transcripts associated with lymphatic endothelial cells.

**Figure 2 supplement 1.**
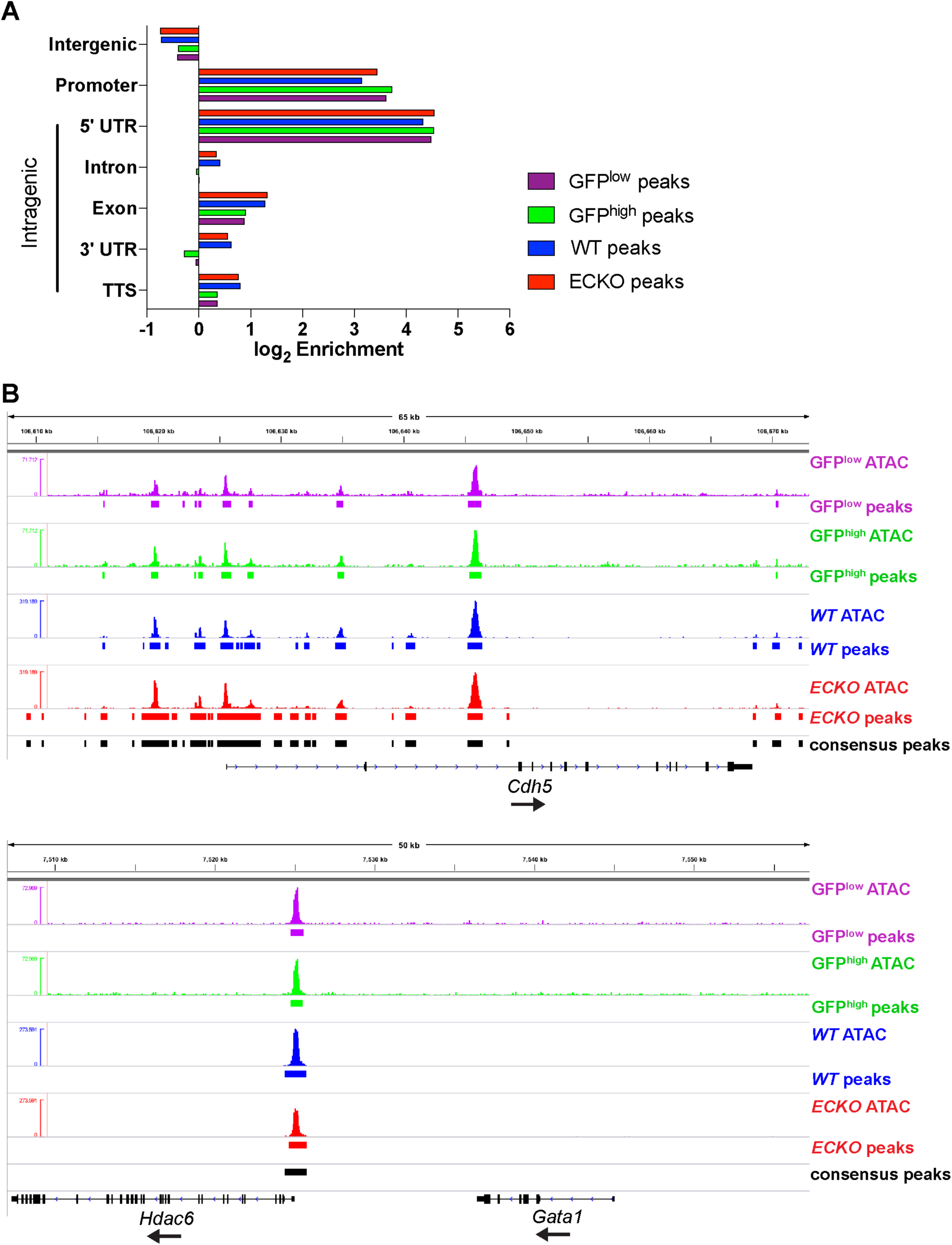
ATAC-seq quality control and peak annotation. (A) Enrichment of GFP^high^, GFP^low^, *S1pr1-ECKO* and *WT* MAECs peaks with respect to distance from UCSC-annotated transcripts (mm9). Promoters are defined as +/- 1 kb from transcription start sites. **(B)** ATAC-seq signal and peaks at *Cdh5* and *Gata1* loci.

**Figure 2 supplement 2.**
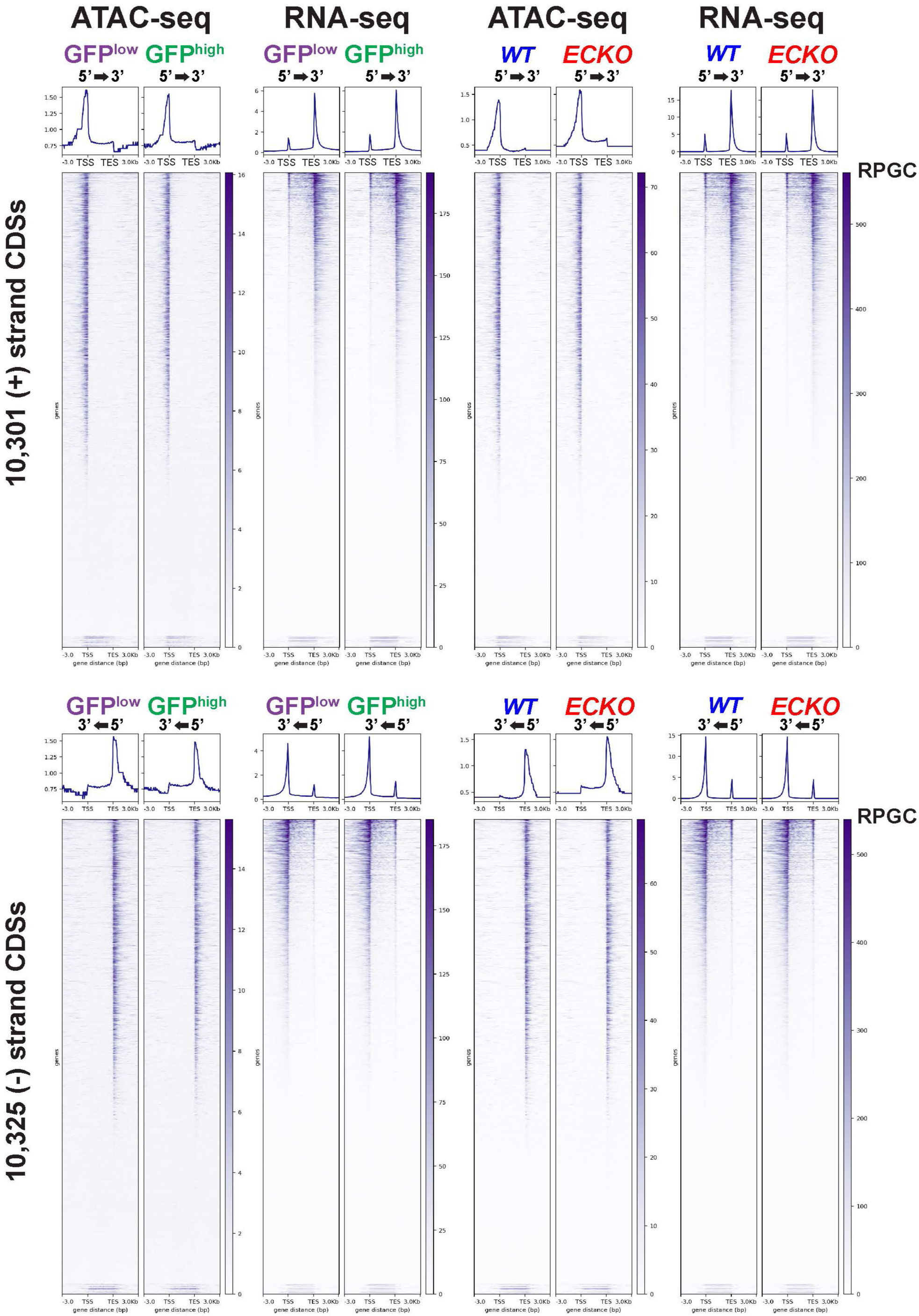
Genome-wide concordance between promoter-proximal chromatin accessibility and mRNA expression in sorted MAEC populations. ATAC-seq and RNA-seq reads (RPGC-normalized) were examined from 3kb upstream and 3kb downstream of all UCSC-annotated CDS’s using the DeepTools ComputeMatrix function. Signal was generated in bin sizes of 50 bp and 100 bp for ATAC-seq and RNA-seq, respectively.

**Figure 2 supplement 3.**
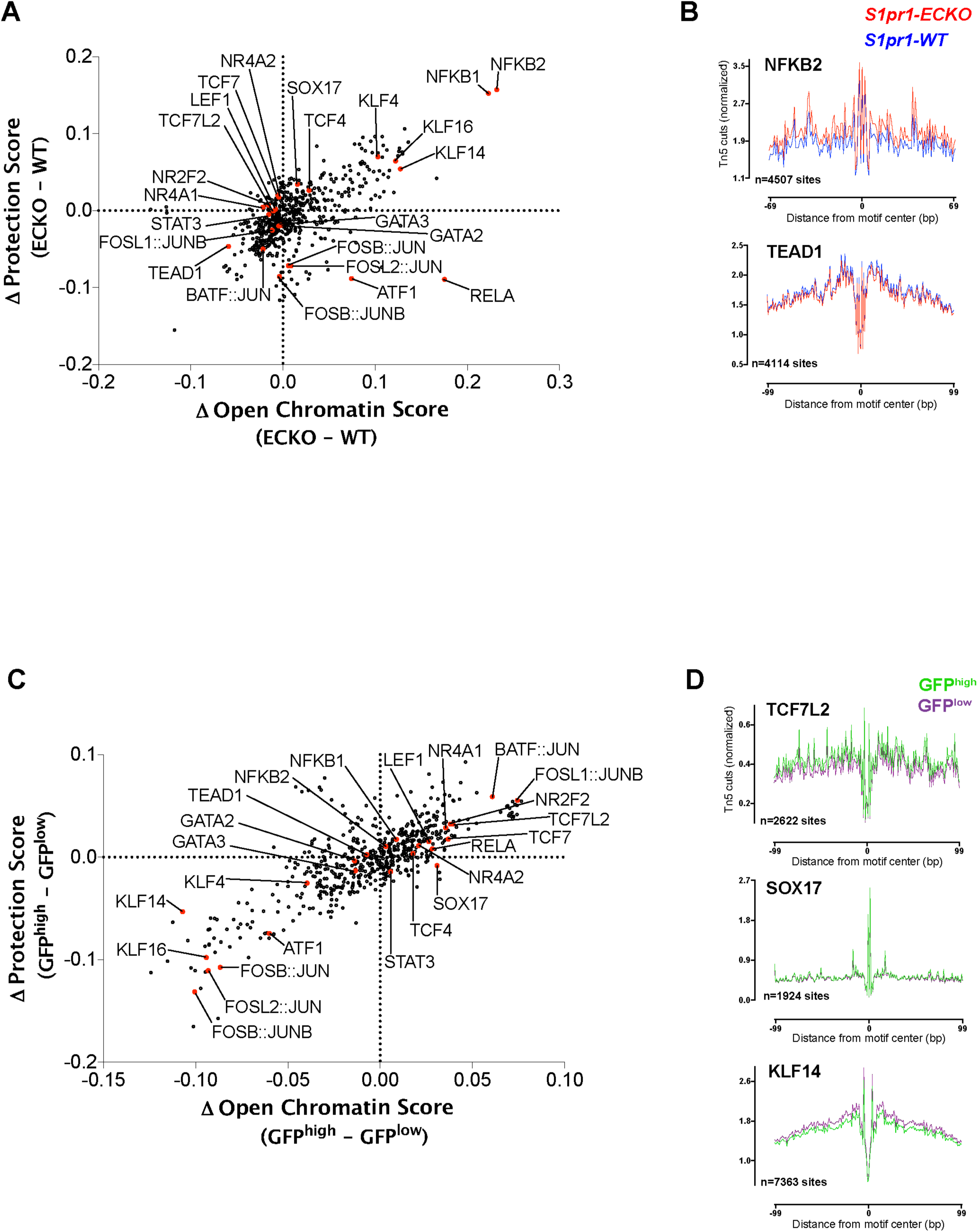
Genome-wide chromatin footprinting with HINT-ATAC. (A) The change in chromatin accessibility scores between GFP^high^ and GFP^low^ MAECs plotted against the change in protection scores for each of the 579 queried JASPAR motifs. **(B)** Lineplots of Tn5 cuts from GFP^high^ and GFP^low^ MAECs averaged over the sites identified in **(A)** for TCF7L2, SOX17, and KLF14 motifs. **(C)** The same plot as **(A)** showing the differences between *S1pr1- ECKO* and *S1pr1-WT* MAECs. **(D)** Lineplots of Tn5 cuts from *S1pr1-ECKO* and *S1pr1-WT* MAECs averaged over the sites identified in **(C)** for NFKB2 and TEAD1 motifs.

**Figure 3 supplement 1.**
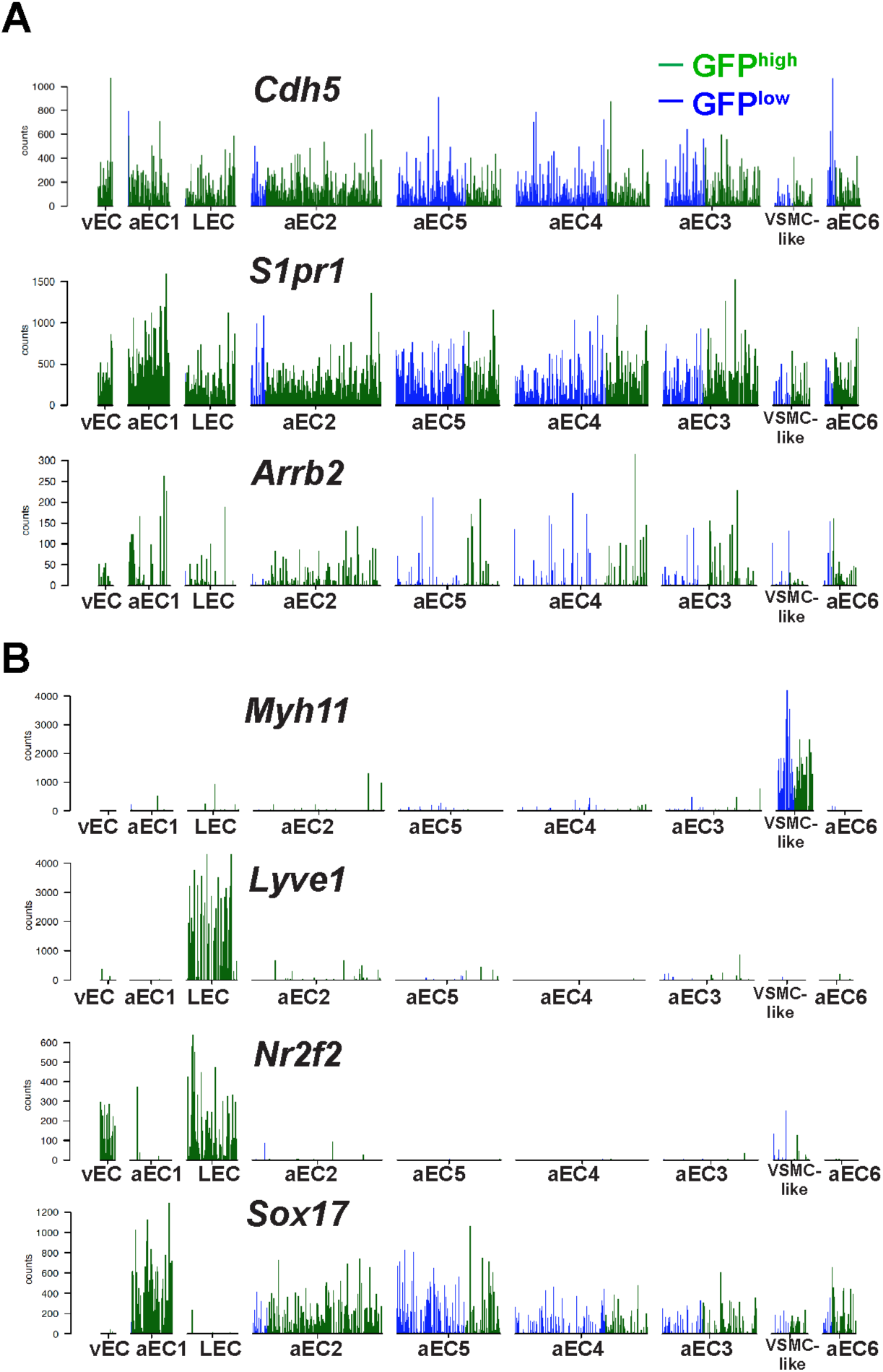
Expression of cluster-defining transcripts in single GFP^high^ and GFP^low^ MAECs. (A-B) Barplots of transcript counts of selected genes in individual GFP^high^ and GFP^low^ MAECs cells. Each line represents a single cell.

**Figure 3 supplement 2.**
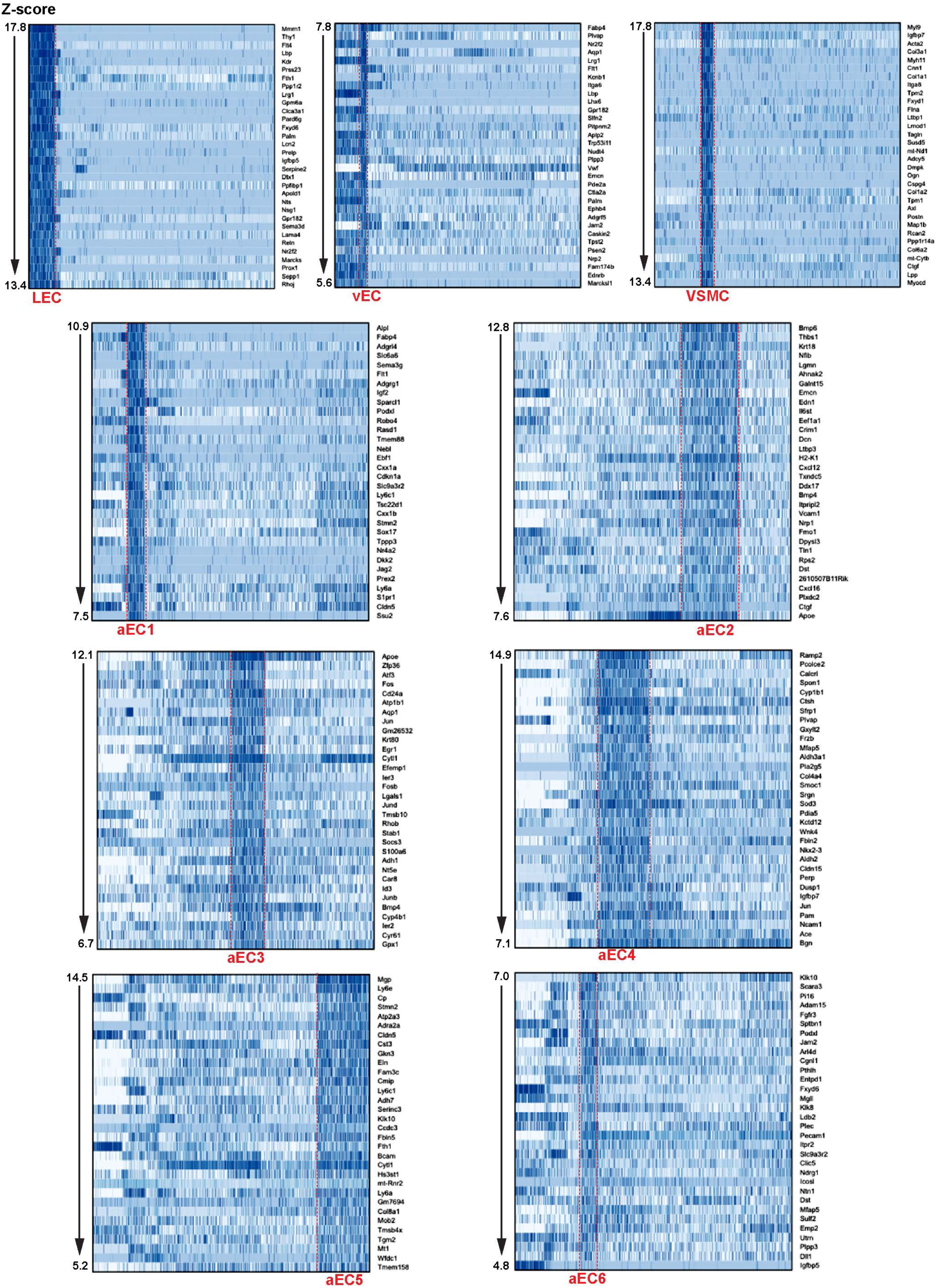
Top markers transcripts for each of the nine clusters defined by scRNA-seq analysis of GFP^high^ and GFP^low^ MAECs. Heatmaps (cells are clustered as shown in Figure 3B) illustrate expression of the top 32 most enriched transcripts in each cluster, according to Z-score.

**Figure 3 supplement 3.**
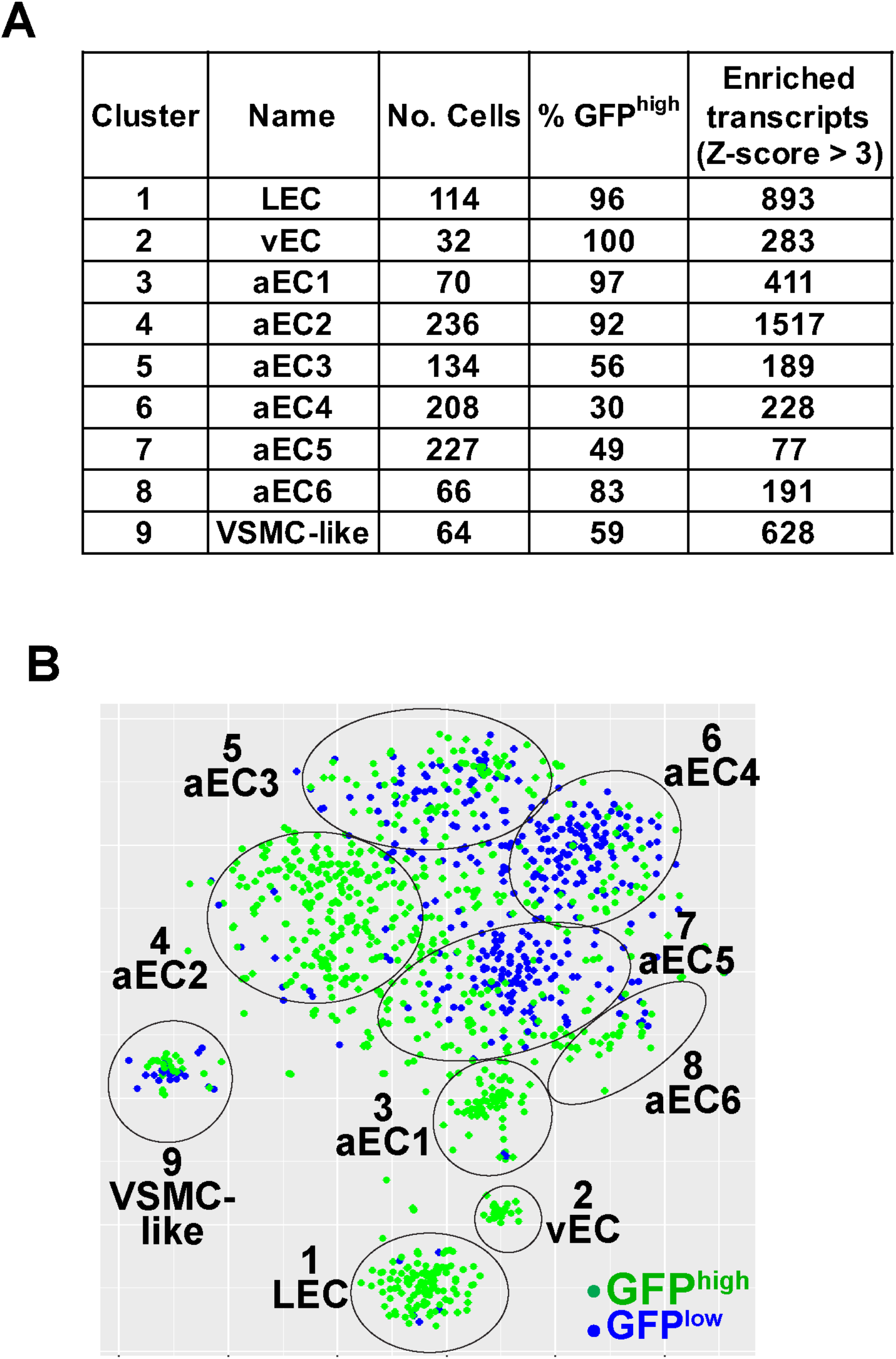
General features and nomenclature of the nine clusters defined by scRNA-seq analysis of GFP^high^ and GFP^low^ MAECs. (A) Table summarizing characteristics of the nine clusters. **(B)** tSNE projection of the embedding shown in Figure 1A with cells color-coded according to GFP expression.

**Figure 3 supplement 4.**
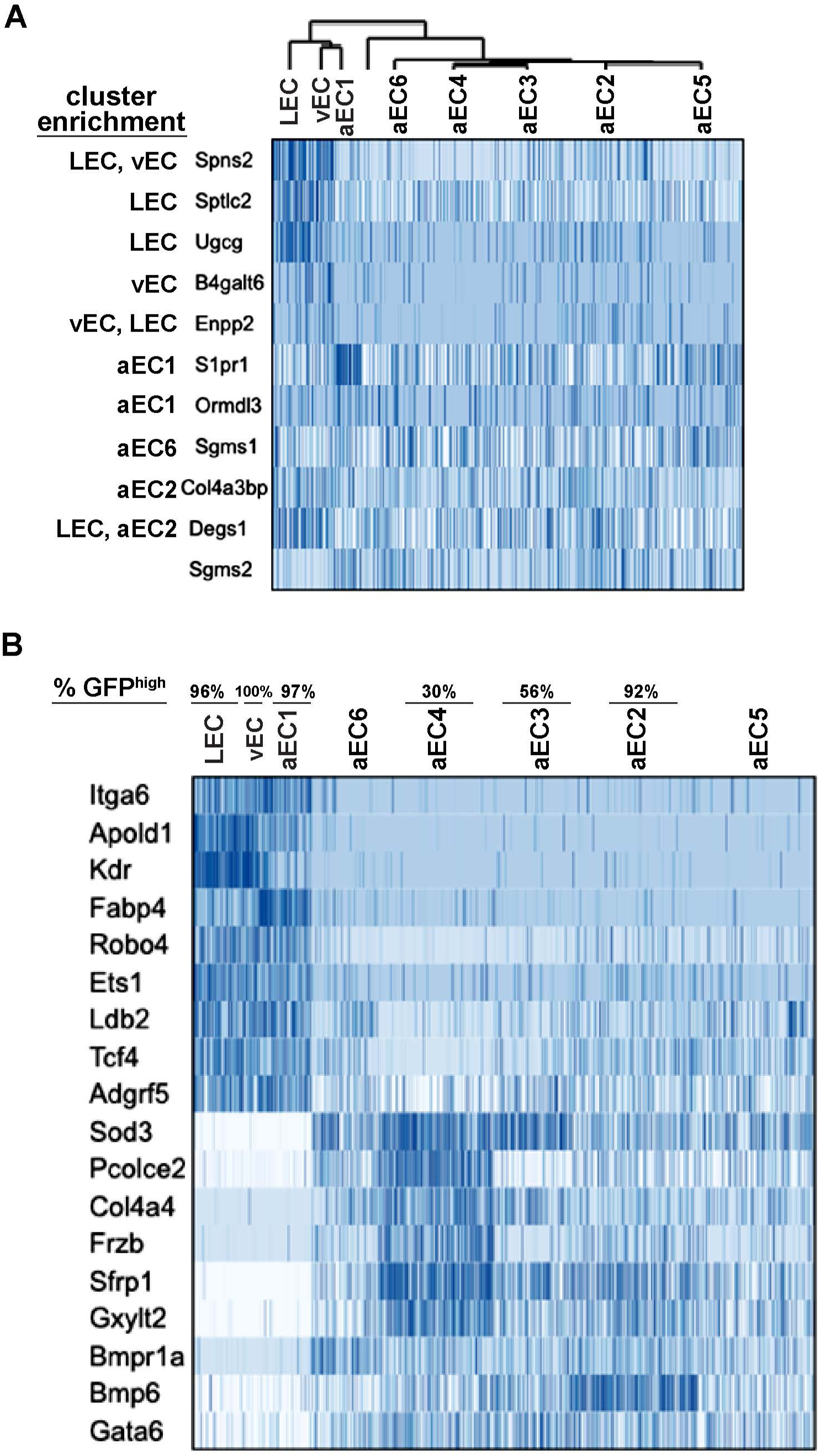
Transcripts co-enriched in LEC, vEC, and aEC1. (A) Heatmaps of sphingolipid-related transcripts showing enrichment in one or multiple EC clusters. **(B)** Heatmaps of examples of transcripts co-enriched and depleted in LEC, vEC, and aEC1 versus aEC2-6. Cell clustering is as shown in Figure 3B.

**Figure 4 supplement 1.**
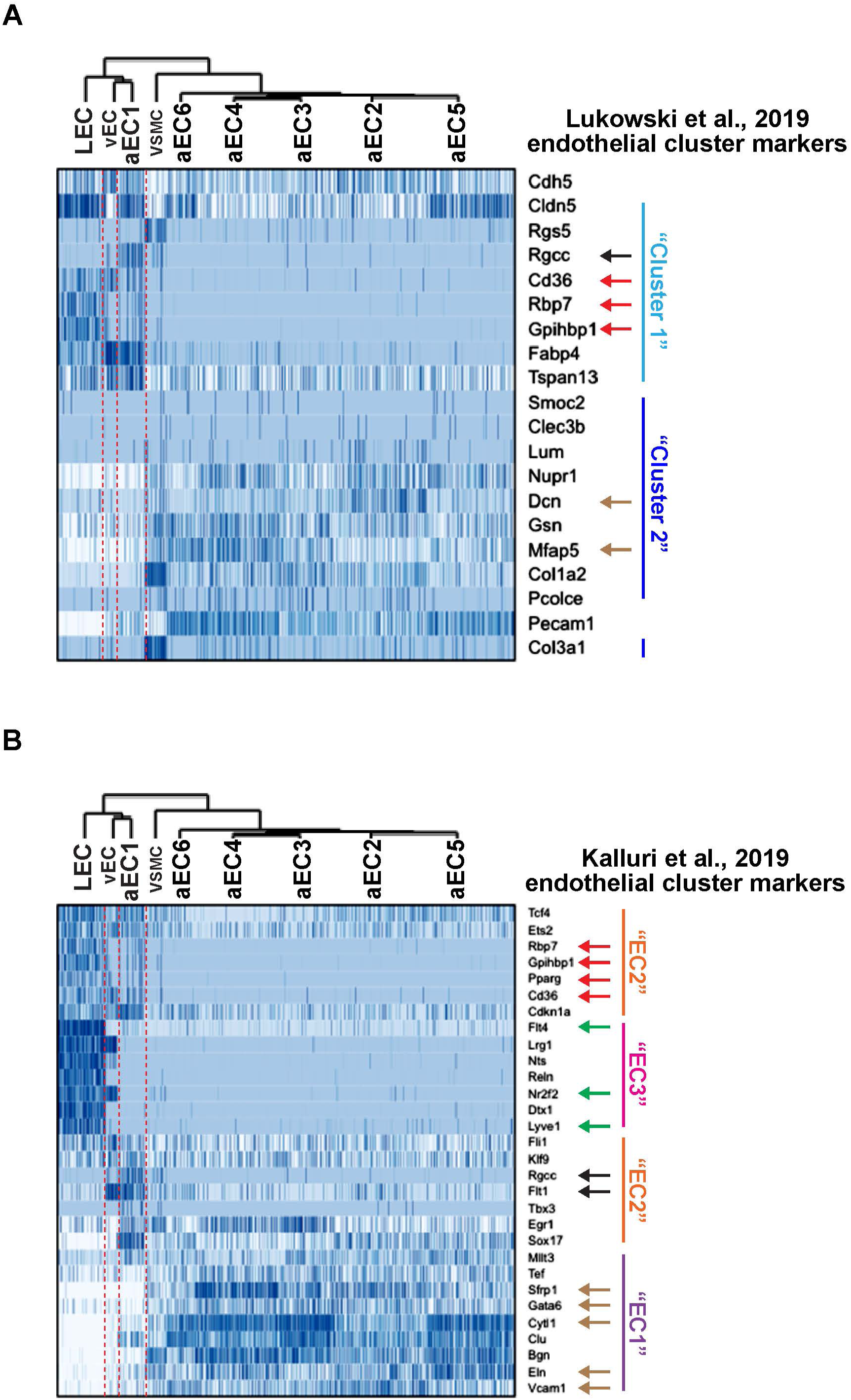
Expression of aortic EC cluster markers from Lukowski et al. (2019) and Kalluri et al. (2019) in LEC, vEC, and aEC1-6. (A) Expression of “Cluster 1” and “Cluster 2” marker genes (Lukowski et al., 2019) in LEC, vEC, and aEC1-6. **(B)** Expression of “EC1”, “EC2”, and “EC3” marker genes (Kalluri et al., 2019) in LEC, vEC, and aEC1-6. Black arrows indicate enrichment in aEC1, red arrows indicate enrichment in both LEC and aEC1, green arrows indicate enrichment in LEC, beige arrows indicate enrichment in aEC2-6 and depletion from LEC, vEC, and aEC1.

**Figure 5 supplement 1.**
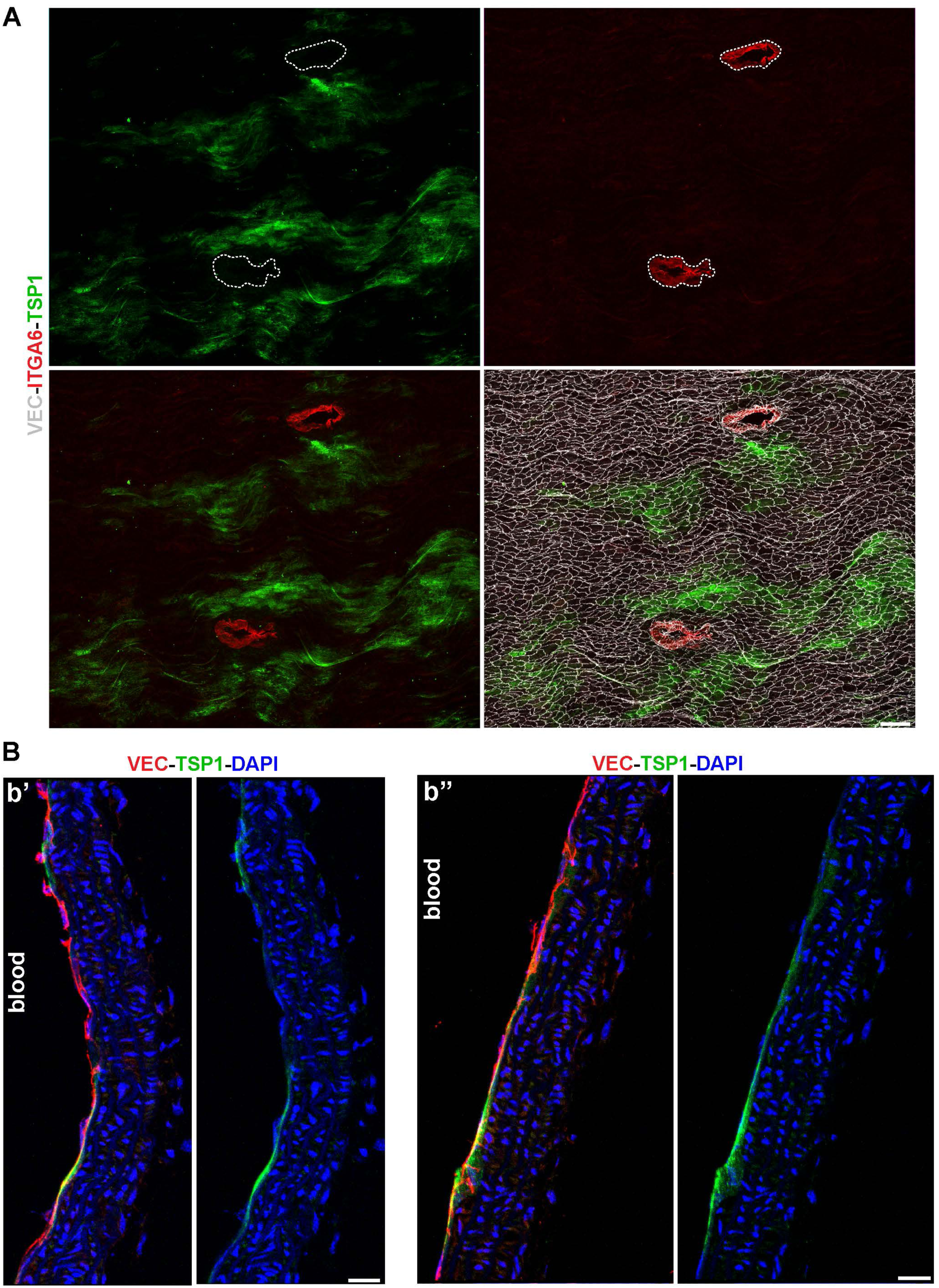
Patchy TSP1 immunoreactivity is endothelial and excluded from thoracic branch point orifices. (A) Whole-mount *en face* preparation of a mouse thoracic aorta immunostained for ITGA6, TSP1, and VEC. Scale bar is 200 µM. **(B)** Two sagittal sections (b’ and b’’) of a mouse thoracic aorta demonstrate TSP1 expression in VEC+ arterial ECs. Sections were counterstained with DAPI to image nuclei. Scale bars are 20 µM.

**Figure 5 supplement 2.**
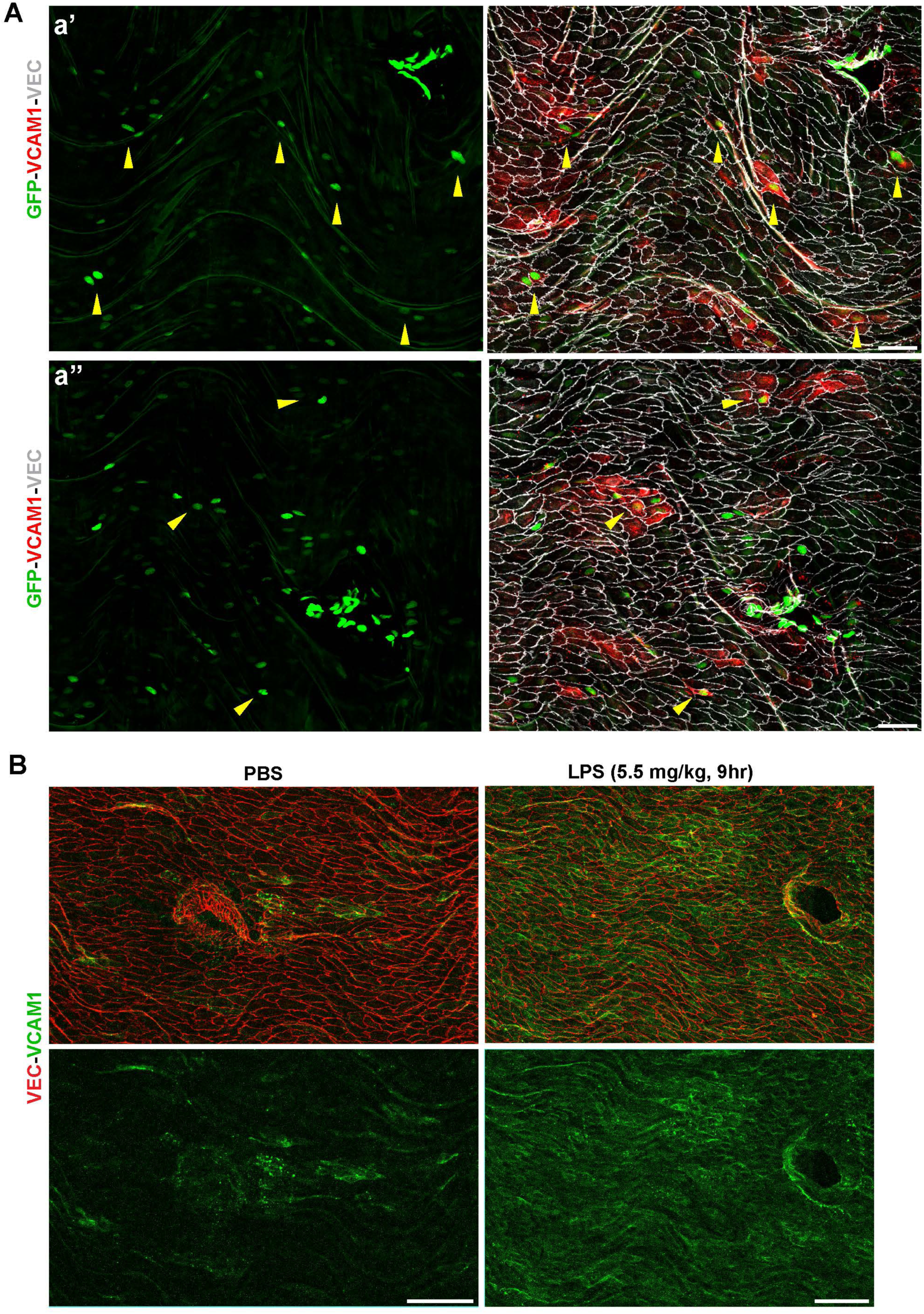
VCAM1 is LPS-inducible and expression is observed in GFP+ ECs distal from thoracic branch point orifices. (A) Two fields (a’ and a’’) from a whole mount *en face* preparation of an S1PR1-GS mouse thoracic aorta immunostained for VCAM1 and VEC. Yellow arrows indicate VCAM1+ GFP+ ECs distal from the branch point orifice. Scale bars are 50 µM. **(B)** Intraperitoneal administration of LPS (5.5 mg/kg, 9hr) results in uniform VCAM1 induction in aortic ECs, visualized by VCAM1 and VEC immunostaining of thoracic aorta whole mount *en face* preparations. Scale bars are 100 µM.

**Figure 7 supplement 1.**
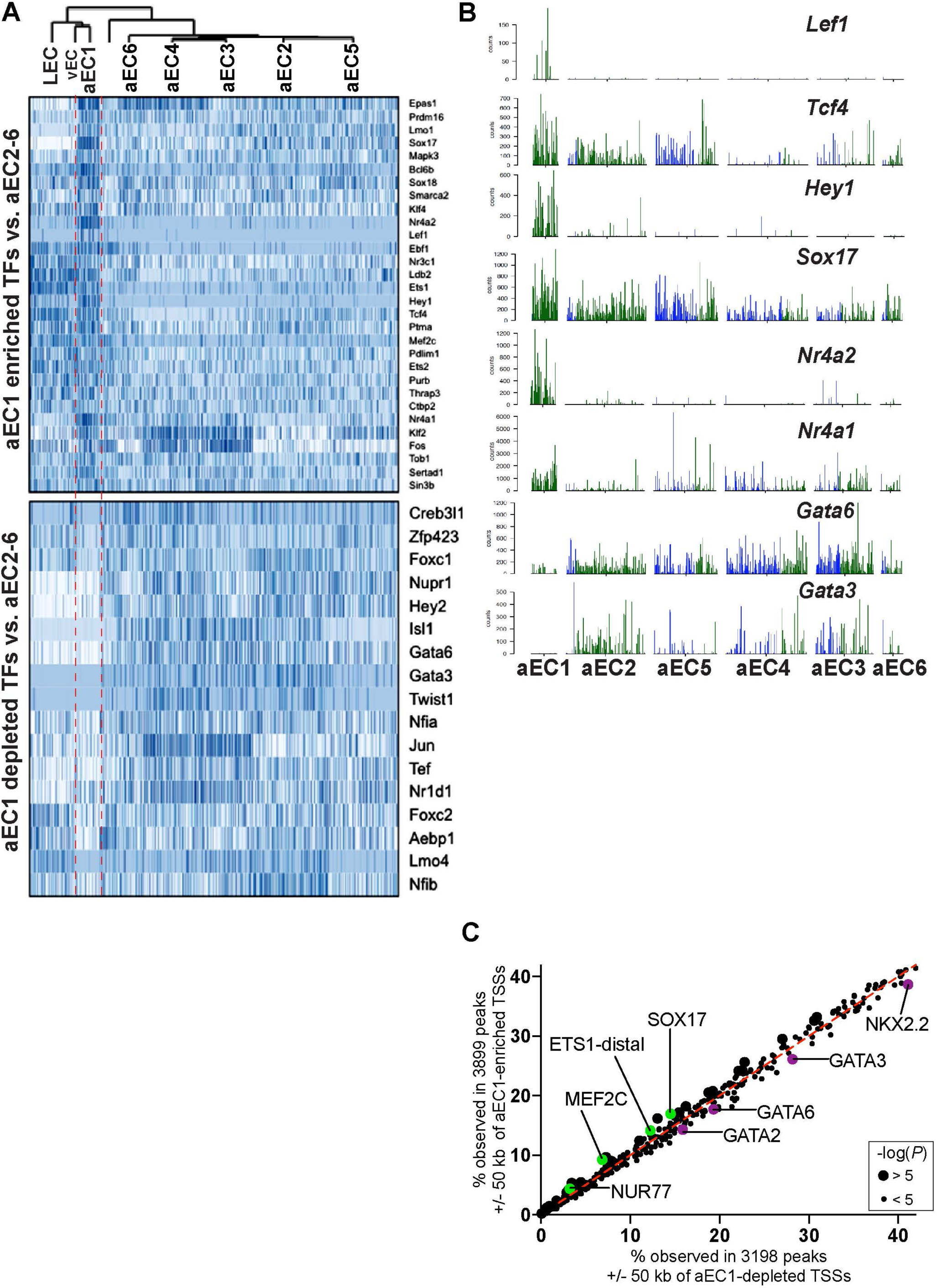
Transcription factors enriched and depleted in aEC1 cells. (A) Heatmaps (cells are clustered as shown in Figure 3B) illustrate expression of transcription factors (TFs) enriched or depleted from aEC1 relative to aEC2-6. **(B)** Barplots illustrate transcript counts of selected TFs from **(A)** in arterial ECs clusters aEC1-6. **(C)** The HOMER script “findmotifsgenome.pl” was used for motif enrichment analysis of merged GFP^high^ and GFP^low^ peaks that intsersected a 100 kb window centered on aEC1-enrichd (Z-score > 3) and depleted (Z-score < -3) TSSs.

**Figure 7 supplement 2.**
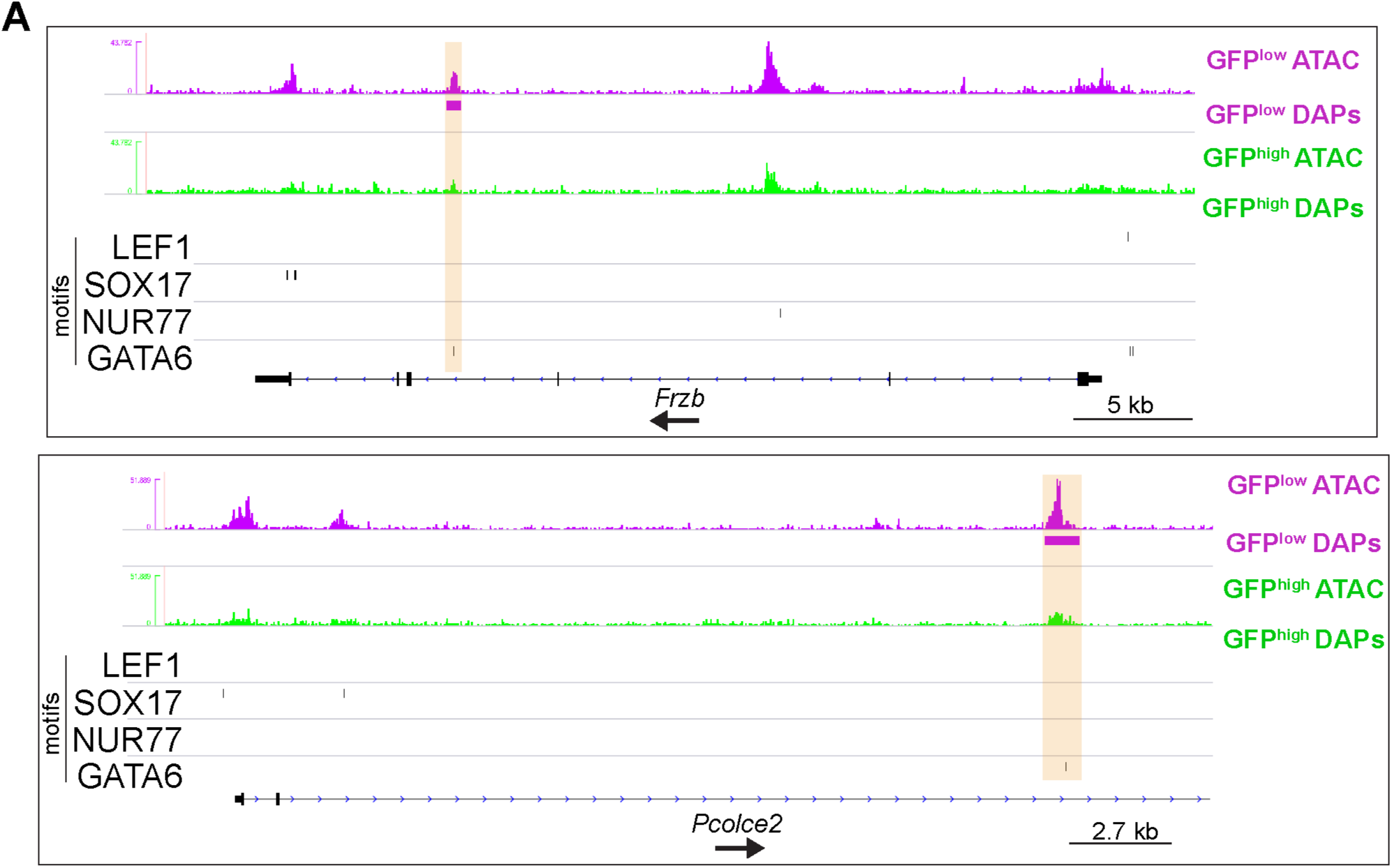
GATA6 motifs in GFP^low^ MAECs open chromatin within aEC4- enriched genes. (A) Genome browser images of *Itga6* and *Gata1*. Peaks with increased accessibility in GFP^low^ MAECs are indicated (GFP^low^ DAPs), as well as LEF1, SOX17, NUR77, and GATA6 motifs in consensus peaks (see Figure 2A). Orange bars highlight co-incidence of GATA6 motifs and increased accessibility in GFP^low^ MAECs.

**Figure 8 supplement 1.**
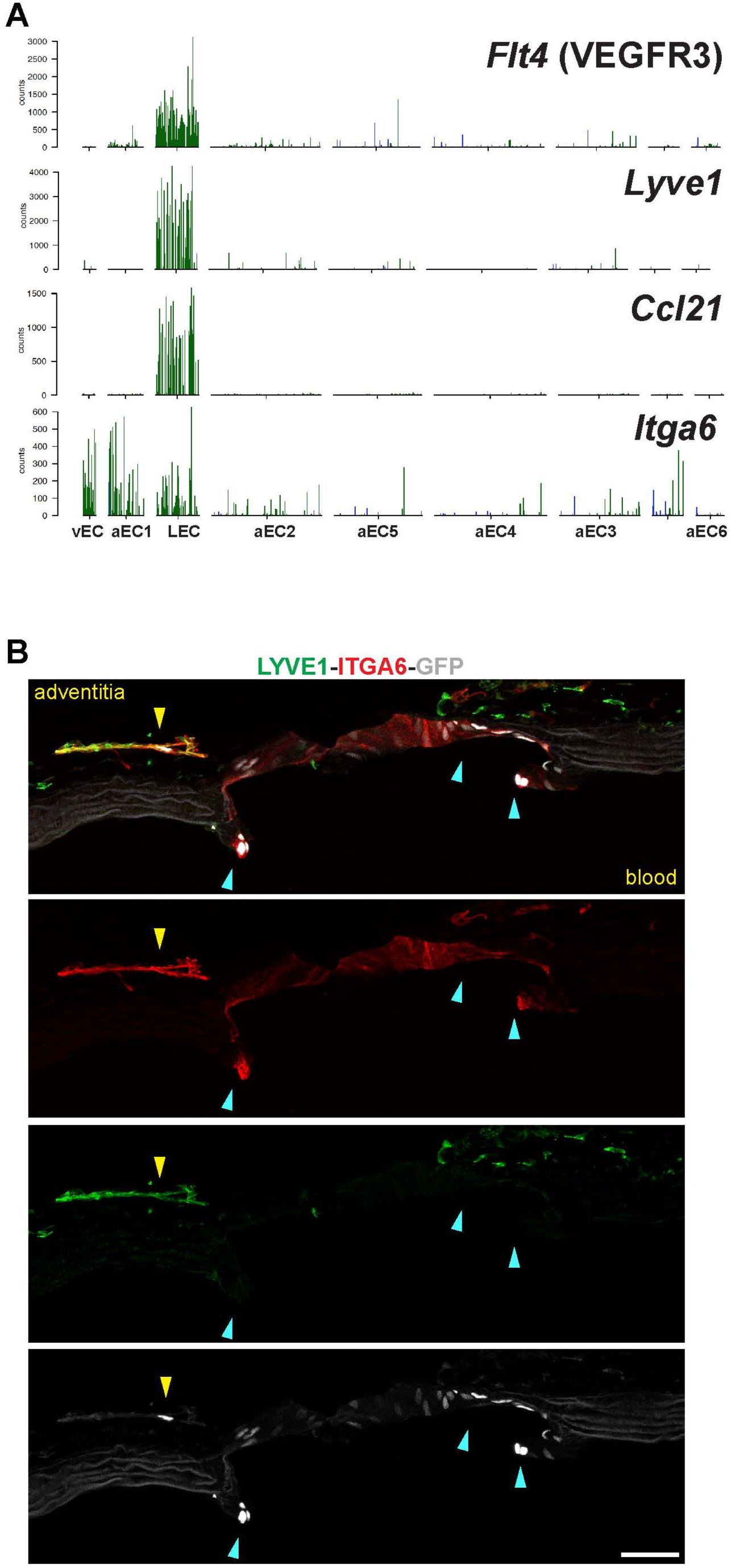
Expression of LEC transcripts and localization of ITGA6+ LEC and branch point arterial EC in close proximity. (A) Barplots illustrate transcript counts of *Flt4* (VEGFR3), *Lyve1*, *Ccl21*, and *Itga6* in each of the nine clusters identified by scRNA-seq analysis. **(B)** Immunostaining of a sagittal section of a S1PR1-GS mouse thoracic aorta for ITGA6 and LYVE1. White arrows indicate GFP+ITGA6+LYVE1- arterial ECs of the branch point orifice. The yellow arrow indicates a GFP+ ECs associated with an adventitial LYVE1+ITGA6+ structure. Scale bar is 50 µM.

**Figure 9 supplement 1.**
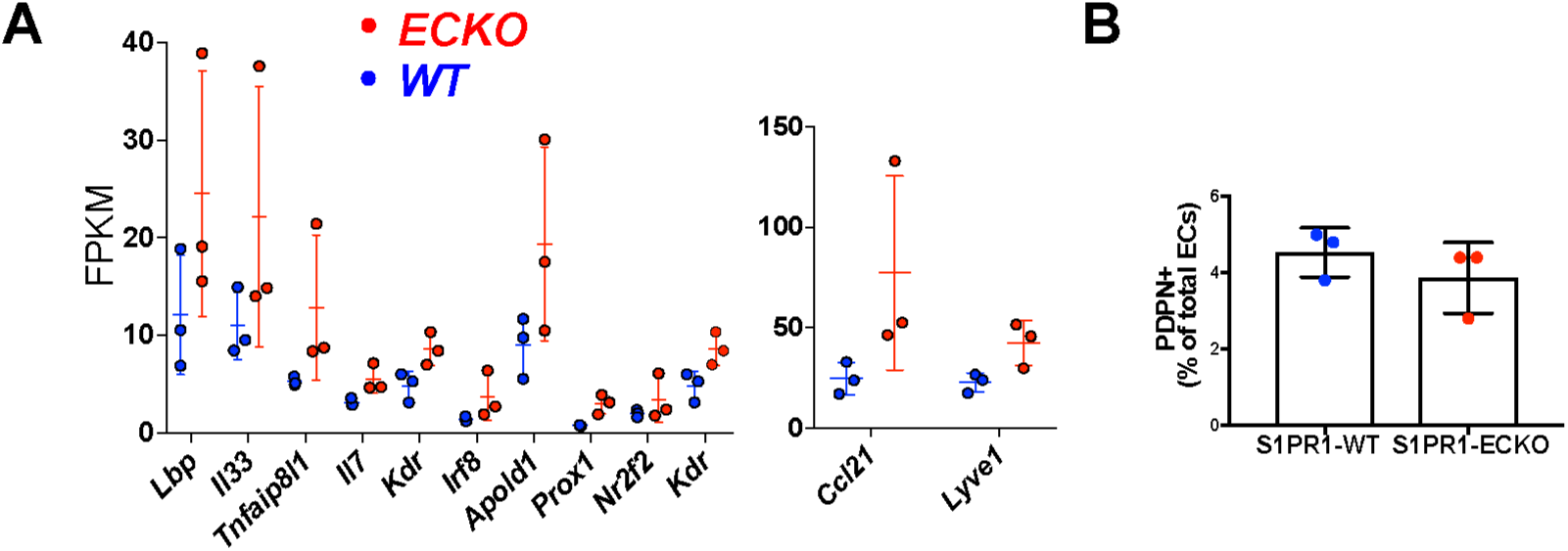
Up-regulation of LEC transcripts in *S1pr1-ECKO* MAECs is not associated with a change in the proportion of aorta-associated PDPN^+^ LECs. (A) FPKMs of selected transcripts from Figure 9B. (B) Aortae were digested and subjected to flow cytometric analysis as described in Figure 1B. PDPN (PE-Cy7-conjugated) was added to the staining cocktail to distinguish aorta-associated LECs from the remainder of aortic ECs.

**Figure 9 supplement 2.**
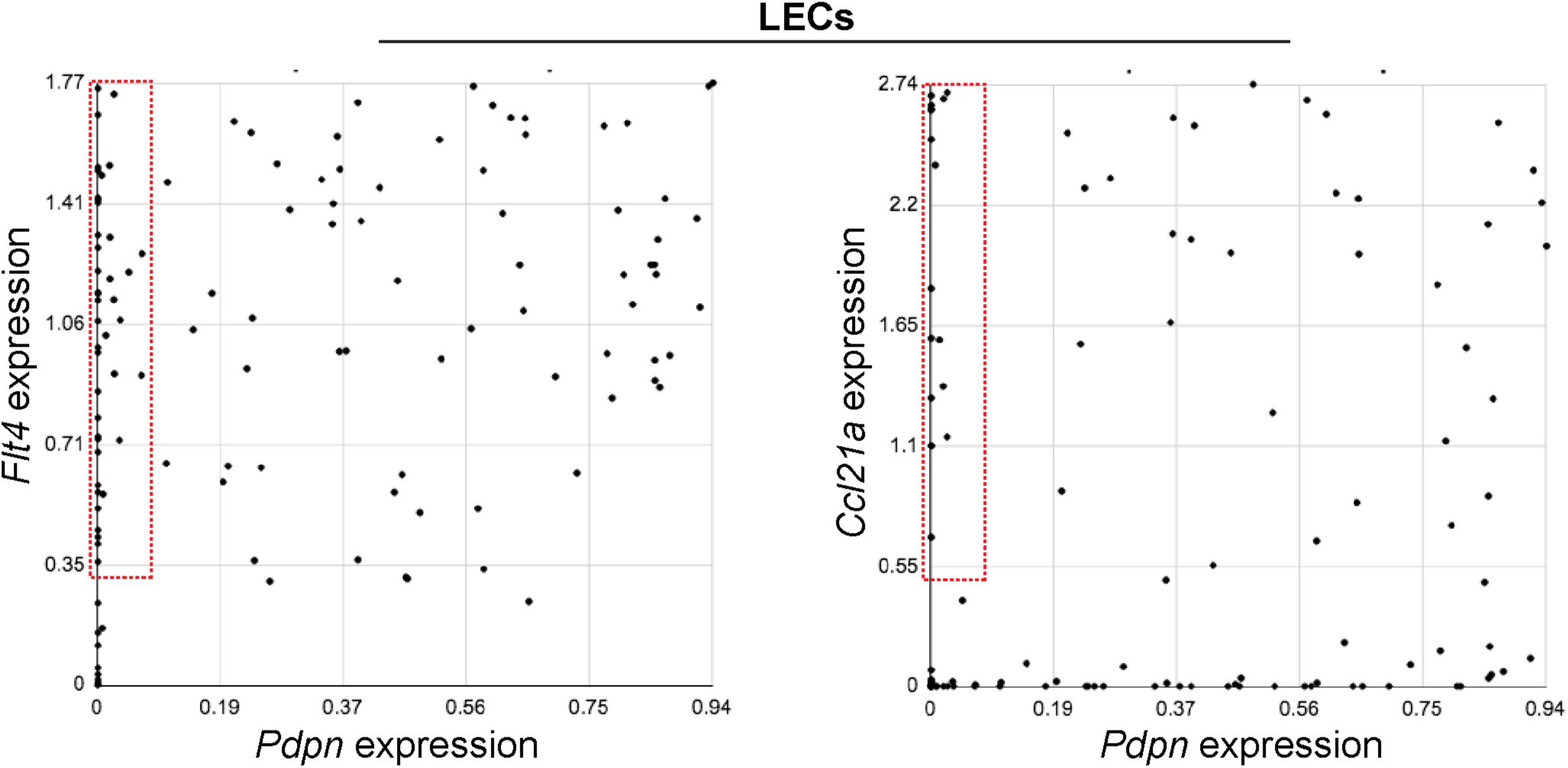
Heterogeneous LEC marker gene expression in aorta-associated LECs. Expression of *Flt4* vs *Pdpn* (left) or *Ccl21a* vs *Pdpn* (right) was compard in individual LECs. The red boxes indicate cell expressing *Flt4* but not *Pdpn* (left) or *Ccl21a* but not *Pdpn* (right).

