## Supplementary File 9 for "Sphingosine 1-phosphate-regulated transcriptomes in heterogenous arterial and lymphatic endothelium of the aorta": Supplementary File 9a Bed.shift.window.10bp.ATAC-seq.sh

```
########################
#Eric Engelbrecht
#bam/bed shift, window
#
#HMS Research Computing
########################

#interactive session, and run without submitting jobs
#the terminal will hang for each step, but you can copy-paste this in all together
bsub -Is -q interactive bash

#"$file" is your filename prefix.  Like "Media_GEN00085494_S1_hg38_merged_sorted.rmdup."
#create the "file" variable like this:
file="Media_GEN00085494_S1_hg38_merged_sorted.rmdup"
#to check what "file" is, 
echo $file

################################################
#samtools: separate forward and reverse strands
################################################

module load seq/samtools/1.3
bsub -q short -W 20 "samtools view -b -F 16 "$file".bam > "$file".forward.bam"
bsub -q short -W 20 "samtools view -b -f 16 "$file".bam > "$file".reverse.bam"

###########################################################
#bamtools: create bed file, pipe and add shift and window
###########################################################

module load seq/BEDtools/2.26.0

#10 base pairs
bedtools bamtobed -i "$file".forward.bam | awk 'BEGIN {OFS="\t"}; {if($6=="+") print $1,$2+4,$2+14,$4,$5,$6; else print $1,$2-5,$2+5,$4,$5,$6}' > "$file".forward.shifted.10.bed
bedtools bamtobed -i "$file".reverse.bam | awk 'BEGIN {OFS="\t"}; {if($6=="+") print $1,$2+4,$2+14,$4,$5,$6; else print $1,$2-5,$2+5,$4,$5,$6}' > "$file".reverse.shifted.10.bed

#1 base pair

bedtools bamtobed -i "$file".forward.bam | awk 'BEGIN {OFS="\t"}; {if($6=="+") print $1,$2+4,$2+4,$4,$5,$6; else print $1,$2-5,$2-5,$4,$5,$6}' > "$file".forward.shifted.1.bed
bedtools bamtobed -i "$file".reverse.bam | awk 'BEGIN {OFS="\t"}; {if($6=="+") print $1,$2+4,$2+4,$4,$5,$6; else print $1,$2-5,$2-5,$4,$5,$6}' > "$file".reverse.shifted.1.bed

#100 base pairs

bedtools bamtobed -i "$file".forward.bam | awk 'BEGIN {OFS="\t"}; {if($6=="+") print $1,$2+4,$2+104,$4,$5,$6; else print $1,$2-5,$2+95,$4,$5,$6}' > "$file".forward.shifted.100.bed
bedtools bamtobed -i "$file".reverse.bam | awk 'BEGIN {OFS="\t"}; {if($6=="+") print $1,$2+4,$2+104,$4,$5,$6; else print $1,$2-5,$2+95,$4,$5,$6}' > "$file".reverse.shifted.100.bed

#####################################
#bamtools: convert a bed back to bam
#####################################

#first download chromosome sizes (aka genome file) from UCSC

/opt/UCSC-tools/bin/fetchChromSizes hg38 > hg38.chrom.sizes

#Reverse strand may truncate, if out of bounds of chrom.sizes
#see "error" file if forward/reverse file sizes don't match up
#nt sequence lost in the bam->bed process, only position retained

bedtools bedtobam -i "$file".forward.shifted.10.bed -g hg38.chrom.sizes > "$file".forward.shifted.10.bam
bedtools bedtobam -i "$file".reverse.shifted.10.bed -g hg38.chrom.sizes > "$file".reverse.shifted.10.bam
```
