## Supplementary File 9 for "Sphingosine 1-phosphate-regulated transcriptomes in heterogenous arterial and lymphatic endothelium of the aorta": Supplementary File 9b STAR.scRNAseq.alignment.sh

```
#!/bin/bash
for i in *.fastq; do STAR --runThreadN 4 --genomeDir /n/groups/shared_databases/igenome/Mus_musculus/UCSC/mm10/Sequence/starIndex --outSAMstrandField intronMotif --twopassMode Basic --readFilesIn $i --outFileNamePrefix ${i%.fastq}; done
```
