## Supplementary File 9 for "Sphingosine 1-phosphate-regulated transcriptomes in heterogenous arterial and lymphatic endothelium of the aorta": Supplementary File 9c Pagoda2.code.make.loom.file.r

```
library(velocyto.R)
library(pagoda2)
library(rhdf5)
read.loom.matrices <- function(file, engine='hdf5r') {
  if (engine == 'h5'){
    cat('reading loom file via h5...\n')
    f <- h5::h5file(file,mode='r');
    cells <- f["col_attrs/CellID"][];
    genes <- f["row_attrs/Gene"][];
    dl <- c(spliced="/layers/spliced",unspliced="/layers/unspliced",ambiguous="/layers/ambiguous");
    if("/layers/spanning" %in% h5::list.datasets(f)) {
      dl <- c(dl,c(spanning="/layers/spanning"))
    }
    dlist <- lapply(dl,function(path) {
      m <- as(f[path][],'dgCMatrix'); rownames(m) <- genes; colnames(m) <- cells; return(m)
    })
    h5::h5close(f)
    return(dlist)
  } else if (engine == 'hdf5r') {
    cat('reading loom file via hdf5r...\n')
    f <- hdf5r::H5File$new(file, mode='r')
    cells <- f[["col_attrs/CellID"]][]
    genes <- f[["row_attrs/Gene"]][]
    dl <- c(spliced="layers/spliced",
            unspliced="layers/unspliced",
            ambiguous="layers/ambiguous")
    if("layers/spanning" %in% hdf5r::list.datasets(f)) {
      dl <- c(dl, c(spanning="layers/spanning"))
    }
    dlist <- lapply(dl, function(path) {
      m <- as(t(f[[path]][,]),'dgCMatrix')
      rownames(m) <- genes; colnames(m) <- cells;
      return(m)
    })
    f$close_all()
    return(dlist)
  }
  else {
    warning('Unknown engine. Use hdf5r or h5 to import loom file.')
    return(list())
  }
}
ldat <- read.loom.matrices("Aorta_GFP_EC.loom")
emat <- ldat$spliced
emat <- emat[,colSums(emat)>=1e3]
rownames(emat) <- make.unique(rownames(emat))
r <- Pagoda2$new(emat,modelType='plain',trim=10,log.scale=T)
r$adjustVariance(plot=T,do.par=T,gam.k=10)
r$calculatePcaReduction(nPcs=100,n.odgenes=3e3,maxit=300)
r$makeKnnGraph(k=30,type='PCA',center=T,distance='cosine');
r$getKnnClusters(method=multilevel.community,type='PCA',name='multilevel')
r$getEmbedding(type='PCA',embeddingType='tSNE',perplexity=50,verbose=T)
par(mfrow=c(1,2))
r$plotEmbedding(type='PCA',embeddingType='tSNE',show.legend=F,mark.clusters=T,min.group.size=10,shuffle.colors=F,mark.cluster.cex=1,alpha=0.3,main='cell clusters')
r$plotEmbedding(type='PCA',embeddingType='tSNE',colors=r$depth,main='depth') 
emat <- ldat$spliced; nmat <- ldat$unspliced; # disregarding spanning reads, as there are too few of them
emat <- emat[,rownames(r$counts)]; nmat <- nmat[,rownames(r$counts)]; # restrict to cells that passed p2 filter
# take cluster labels
cluster.label <- r$clusters$PCA$multilevel # take the cluster factor that was calculated by p2
cell.colors <- pagoda2:::fac2col(cluster.label)
# take embedding form p2
emb <- r$embeddings$PCA$tSNE
cell.dist <- as.dist(1-armaCor(t(r$reductions$PCA)))
emat <- filter.genes.by.cluster.expression(emat,cluster.label,min.max.cluster.average = 0.2)
nmat <- filter.genes.by.cluster.expression(nmat,cluster.label,min.max.cluster.average = 0.05)
length(intersect(rownames(emat),rownames(nmat)))
fit.quantile <- 0.02
rvel.cd <- gene.relative.velocity.estimates(emat,nmat,deltaT=1,kCells=25,cell.dist=cell.dist,fit.quantile=fit.quantile, n.cores=2)
show.velocity.on.embedding.cor(emb,rvel.cd,n=200,scale='sqrt',cell.colors=ac(cell.colors,alpha=0.5),cex=0.8,arrow.scale=3,show.grid.flow=TRUE,min.grid.cell.mass=0.5,grid.n=40,arrow.lwd=1,do.par=F,cell.border.alpha = 0.1, n.cores=2)
dev.off()
save(emat, file="emat.GFPonly.RData")
save(nmat, file="nmat.GFPonly.RData")
save(rvel.cd, file="rvel.cd.GFPonly.RData")
save(emb, file="emb.GFPonly.RData")
save(r, file="r.GFPonly.RData")
q()
y
```
